# Volatility estimates increase choice switching and relate to prefrontal activity in schizophrenia

**DOI:** 10.1101/227967

**Authors:** L. Deserno, R. Boehme, C. Mathys, T. Katthagen, J. Kaminski, K. E. Stephan, A. Heinz, F. Schlagenhauf

## Abstract

**Background:** Reward-based decision-making is impaired in patients with schizophrenia (PSZ) as reflected by increased choice switching. The underlying cognitive and motivational processes as well as associated neural signatures remain unknown. Reinforcement Learning (RL) and hierarchical Bayesian learning account for choice switching in different ways. We hypothesized that enhanced choice switching, as seen in PSZ during reward-based decision-making, relates to higher-order beliefs about environmental volatility and examined the associated neural activity.

**Methods:** 46 medicated PSZ and 43 healthy controls (HC) performed a reward-based decision-making task requiring flexible responses to changing action-outcome contingencies during functional Magnetic Resonance Imaging (fMRI). Detailed computational modeling of choice data was performed, including RL and the hierarchical Gaussian filter (HGF). Trajectories of learning from computational modeling informed the analysis of fMRI data.

**Results:** A three-level HGF accounted best for the observed choice data. This model revealed a heightened initial belief about environmental volatility and a stronger influence of volatility on lower-level learning of action-outcome contingencies in PSZ as compared to HC. This was replicated in an independent sample of non-medicated PSZ. Beliefs about environmental volatility were reflected by higher activity in dorsolateral prefrontal cortex of PSZ as compared to HC.

**Conclusions:** Our study suggests that PSZ inferred the environment as overly volatile, which may explain increased choice switching. In PSZ, activity in dorsolateral prefrontal cortex was more strongly related to beliefs about environmental volatility. Our computational phenotyping approach may provide useful information to dissect clinical heterogeneity and could improve prediction of outcome.

## Introduction

Cognitive and motivational deficits are important characteristics of patients with schizophrenia (PSZ) associated with clinical and social outcome^1–5^. Reward-based learning and decision-making require the integration of cognition and motivation and are impaired in PSZ^6, 7^. These impairments are present at the onset of the disorder, are independent of lower general IQ, remain stable over time^8, 9^ and have been proposed as neurocognitive markers with potential clinical utility^10^. However, the mechanisms and associated neural signatures remain to be identified.

Flexible reward-based learning and decision-making can be probed via variants of reversal learning (e.g.^11^). In such tasks, PSZ show increased switching between choice options^8, 12–17^. The mechanisms of this instable behavior remain unknown but can be targeted by computational modeling^18^. In Reinforcement Learning (RL^19^), choices are selected based on expected values, which are learned by weighting reward prediction errors (RPEs) with a learning rate. RPEs closely align with phasic dopamine^20, 21^. Considering enhanced presynaptic dopamine synthesis capacity in striatum of PSZ^22, 23^, this could translate into enhanced phasic dopamine in PSZ, which in turn might result in increased learning rates^24^. This could theoretically account for instable behavior in PSZ, but increased learning rates were not found (for review see ^18, 24, 25^).

Theories of predictive coding^26^ and hierarchical Bayesian inference hypothesize that symptoms of PSZ^27–29^ are a consequence of false inference about the world due to altered precision attributed to beliefs at different hierarchical levels. Dysfunction at higher levels, which are thought to extract and represent general and stable features of the environment, might lead to experiencing the world as being more or less volatile. With regard to positive symptoms^30^, this is supported by empirical evidence (e.g.^31^). When applying this framework to reward-based decision-making, beliefs about the probability of rewards are formed at lower levels but are also determined by learning about the volatility of reward probabilities^32^. This environmental volatility is related to learning from lower-level RPEs in that it scales the belief update. Thus, a belief in high environmental volatility can induce rapid updates of lower-level beliefs about reward probabilities and promote enhanced choice switching in PSZ.

Striatal and prefrontal activity is reduced during reward anticipation and receipt in PSZ^33–35^. Reduced striatal RPE activity was observed in non-medicated^17^ but not in medicated PSZ^15, 36^. Neural correlates of hierarchical Bayesian learning were demonstrated in functional Magnetic Resonance Imaging (fMRI) studies in healthy individuals^37, 38^ linking volatility and uncertainty to activity in fronto-striatal circuits^32, 39^. While neural correlates of hierarchical Bayesian learning were successfully used to distinguish between individuals with and without hallucinations and PSZ with and without psychosis^31^, this has not yet facilitated an understanding of the cognitive and motivational processes underlying impairments in flexible reward-based decision-making.

Here, we used a reward-based reversal-learning task during fMRI in PSZ and healthy controls (HC). Computational modeling was applied to the behavioral data by comparing RL and a hierarchical Bayesian learning model, the Hierarchical Gaussian Filter (HGF)^40, 41^. We hypothesized that enhanced choice switching in PSZ relates to higher-order beliefs about the volatility of the environment and examined the associated neural activity as measured by fMRI.

## Materials and Methods

### Participants and instruments

46 medicated PSZ and 43 HC were included (see S-Table 1). Measures used to characterize participants are summarized in S-Table 1 and supplementary material. Written informed consent was obtained from all participants. The study was performed in accordance with the Declaration of Helsinki and approved by the local ethics committee of Charité Universitätsmedizin.

### Task

Participants performed a task requiring flexible decision-making during fMRI^42–44^. The task had 160 trials, each with a choice between two cards (Figure 1A). The selected card resulted in a monetary win or a monetary loss of 10 Eurocents. One card was initially assigned with a reward probability of 80% and a loss probability of 20% (vice versa for the other card). The task had a simple higher-order structure (Figure 1B): an anti-correlation between the reward probabilities; whenever one card was associated with a probability of 80%, the other card would be associated with a probability of 20%. Reward contingencies were stable for the first 55 trials (‘pre-reversal’) and for the last 35 trials (‘post-reversal’). During the ‘reversal’ phase, contingencies changed four times, after 15 or 20 trials, respectively. For more details, see supplementary material.

**Figure 1.**
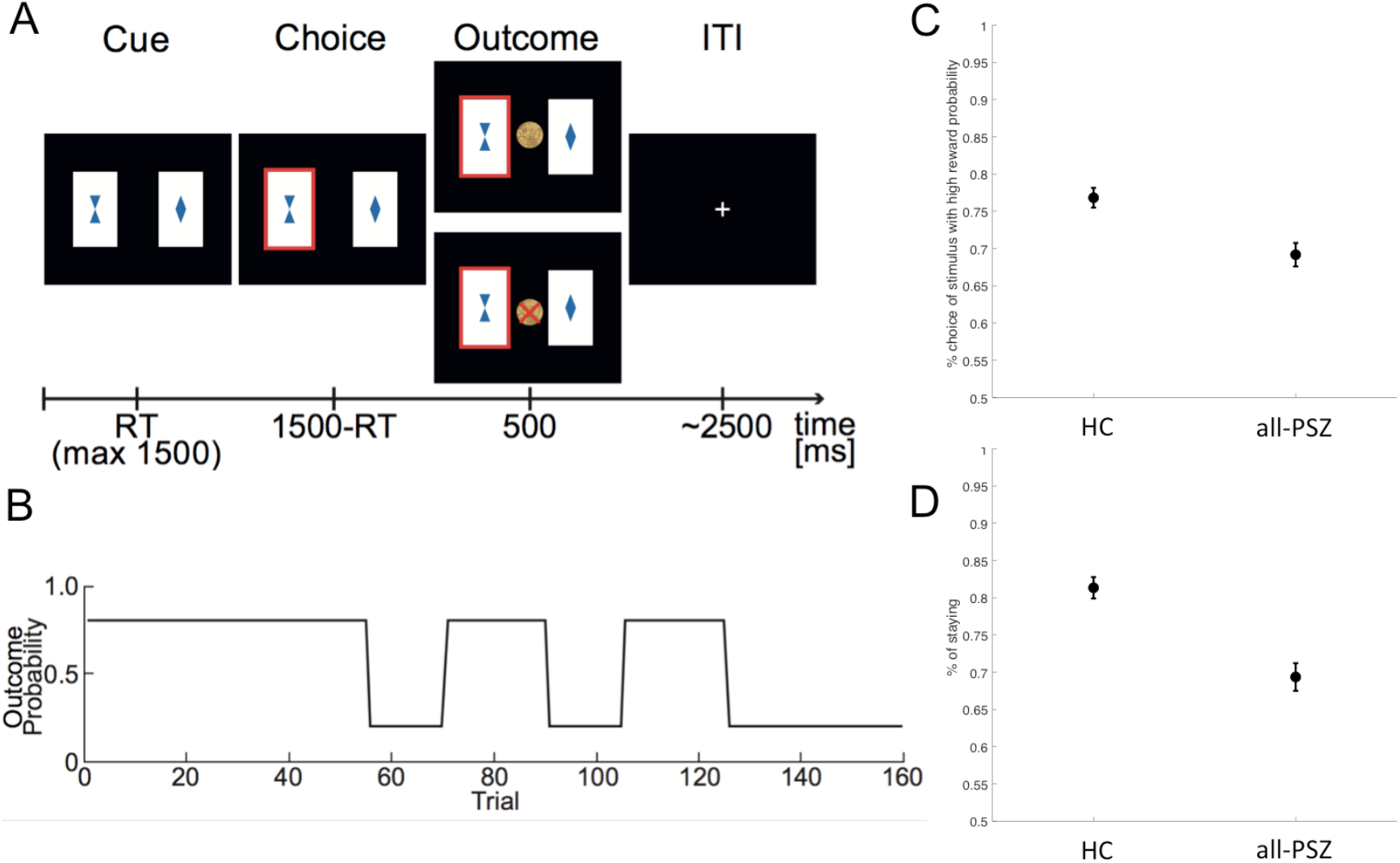
(A) Trial sequence from the decision-making task. (B) Reward probabilities of both choice options were perfectly anti-correlated and were stable for the first 55 trials (‘pre-reversal’), changed four times, after 15 or 20 trials, in the ‘reversal’ phase and remained stable for the last 35 trials (‘post-reversal’). (C) Percent choices of the stimulus with 80% reward probability was significantly lower in the PSZ group (main effect of group F=14.52, p<0.001). (D) PSZ were less likely to repeat the previous action independent of feedback received in the previous trial (main effect of group F=27.77, p<0.001, feedback x group interaction F=0.02, p=0.89).

### Analysis of choice behavior

Performance was quantified by ‘correct’ choices of the stimuli with high (80%) reward probability and analyzed using repeated-measures ANOVA with the between-subject factor ‘group’ and the within-subject factor ‘phase’ (pre-reversal, reversal, post-reversal). Repeated-measures ANOVA was used to test the effect of feedback on subsequent choices (‘win-stay’ and ‘lose-stay’).

### Computational models of learning

In RL, the difference between received rewards and expectations, the RPE, is used to update expectations for the chosen stimulus (weighted by the learning rate *α*). For comparison with previous work^17, 42–44^, we included RL with separate learning rates for reward and loss trials (RL1, RL2).

The HGF describes learning as a process of inductive inference under uncertainty. It considers hierarchically organized states, in which learning at a higher-level state determines learning at a lower-level state by dynamically adjusting the lower level’s learning rate. In our case, the top level represents environmental volatility (how likely a change in action-outcome contingencies is to occur). This top-level estimate is dynamically coupled with learning at the lower level (see Figure 2). Trial-by-trial updates of posterior means at each level are proportional to the prediction error (PE) from the level below weighted by a precision ratio. See supplementary material for equations. We were particularly interested in environmental volatility (*µ*_3_) and its coupling with the lower level (κ) and thus inferred subject-specific parameters. We included a two-level variant (HGF2) to test whether the representation of volatility in the three-level HGF3 made it superior in explaining behavior.

**Figure 2.**
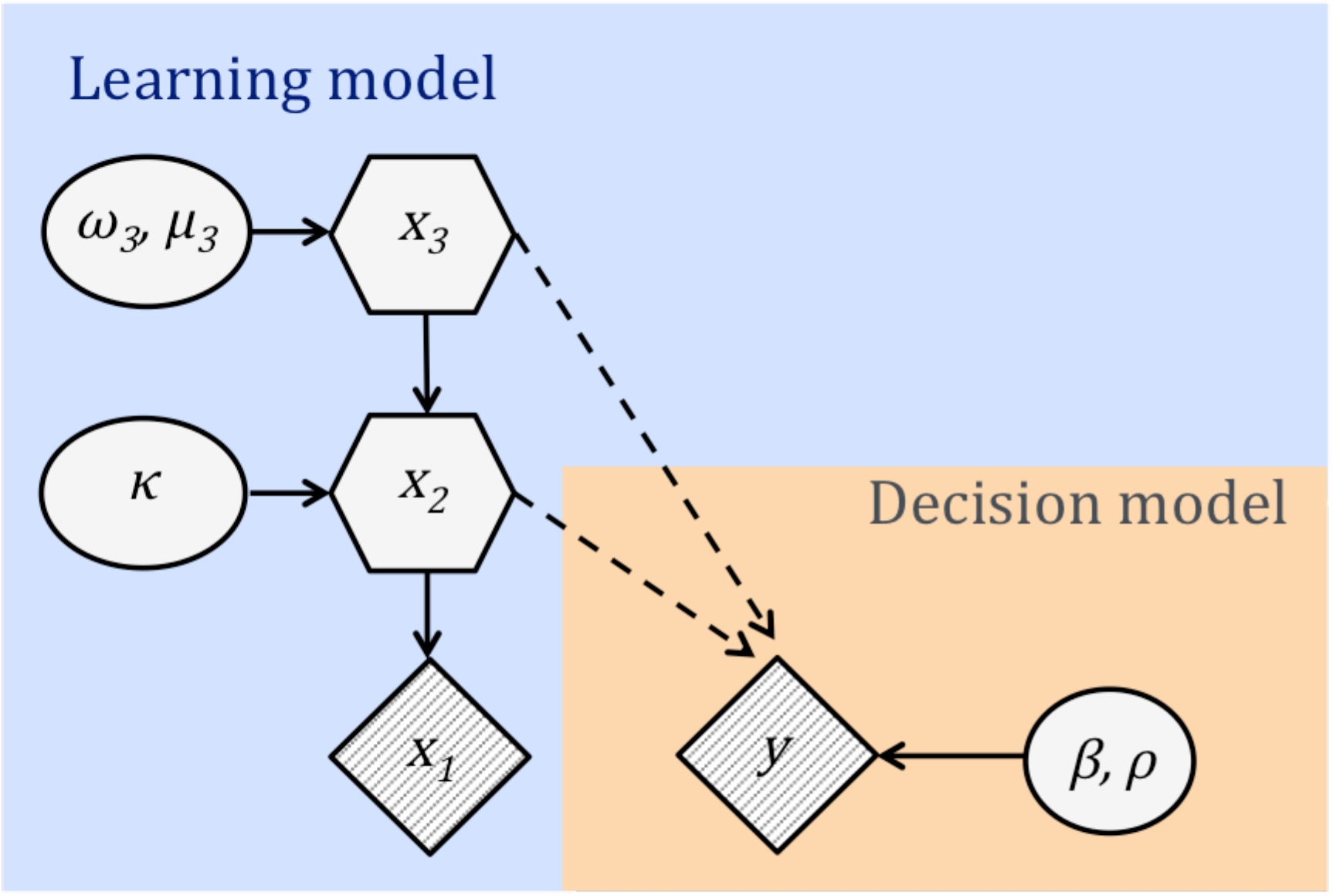
Model graph. The HGF deploys hierarchically organized states, in which learning about environmental volatility at a higher-level state *x_3_* determines lower-level learning about reward probabilities *x_2_*. The lowest level, x1, is binary and corresponds to a choice being rewarded (*x_1_*=1) or not (*x_1_*=0) at a given trial. The probability of a choice being rewarded is a logistic sigmoid function of *x_2_*: *p*(*x_1_*=1) = *s*(*x_2_*). y represents the response of the subject. Shaded quantities are observed. Solid lines indicate dependence in the generative model. Dashed lines indicate dependence on inferred quantity (the generative model for *y* depends on *μ_2_* and *μ_3_*, the inferred values of *x_2_* and *x_3_*). The constant step size ω_3_ is the evolution rate of environmental volatility. κ reflects the coupling between the levels. The best-fitting model was a three-level implementation (HGF3-DU-V) with double-updating (not illustrated) together with a decision model capturing choice repetition separately after rewards and losses (ρ), third-level environmental volatility determining decision noise, and the initial belief about environmental volatility *µ_3_* as an additional parameter inferred from the data.

HGF and RL provide different ways to learn expectations about rewards and both update expectations of the chosen card only (“single-update”, SU). Based on the anti-correlated task structure, we implemented a variant of each model updating values (RL) or posterior means (HGF) of the unchosen card simultaneously, i.e. an increase of the chosen card implies a decrease of the unchosen card (“double-update”, DU). For equations, see supplementary material. SU and DU variants of each model (RL1, RL2, HGF2, HGF3) were fit to the choice data (Table 1).

**Table 1.** Bayesian Model Selection (BMS). BMS was governed by protected exceedance probabilities (PXP) to protect against the risk that differences in model evidences occur by chance. In this table we also report exceedance probabilities (PX) and expected posterior probabilities (PP), also compare supplementary material. PX describes the probability of a model to exceed all other models in the comparison set, the probability that expected PPs differ; HC = healthy controls, PSZ = patients with schizophrenia, RL = Reinforcement Learning with one learning rate (1) or separate learning rates for rewards and losses (2). HGFX = Hierarchical Gaussian Filter with 2 or 3 levels, SU = single update, DU = double update, HGF3-**-V = three-level HGF with environmental volatility linked to decision noise with either SU or DU.

|  |  | RL1-SU | RL1-DU | RL2-SU | RL2-DU | HGF2-SU | HGF2-DU | HGF3-SU | HGF3-DU | HGF3-SU-V | HGF3-DU-V |
| --- | --- | --- | --- | --- | --- | --- | --- | --- | --- | --- | --- |
| **HC+PSZ**<br>*all n=89* | *PP* | 9.4 | 5.1 | 7.4 | 3.0 | 3.1 | 7.0 | 3.3 | 7.7 | 5.3 | 48.8 |
|  | *XP* | 0 | 0 | 0 | 0 | 0 | 0 | 0 | 0 | 0 | 100 |
|  | *PXP* | 0 | 0 | 0 | 0 | 0 | 0 | 0 | 0 | 0 | **99.9** |
| **HC**<br>*all n=43* | *PP* | 4.4 | 4.7 | 4.4 | 3.8 | 2.7 | 5.1 | 2.7 | 5.3 | 5.8 | 61.1 |
|  | *XP* | 0 | 0 | 0 | 0 | 0 | 0 | 0 | 0 | 0 | 100 |
|  | *PXP* | 0 | 0 | 0 | 0 | 0 | 0 | 0 | 0 | 0 | **100** |
| **PSZ**<br>*all n=46* | *PP* | 11.8 | 7.1 | 11.6 | 4.75 | 8.3 | 10.0 | 9.3 | 10.9 | 4.9 | 21.4 |
|  | *XP* | 6.9 | .7 | 6.21 | .1 | 1.4 | 3.4 | 2.5 | 4.6 | .2 | 74.0 |
|  | *PXP* | 10.0 | 10.0 | 10.0 | 10.0 | 10.0 | 10.0 | 10.0 | 10.0 | 10.0 | **10.1** |
| **HC+PSZ**<br>*fit n=73* | *PP* | 9.4 | 5.1 | 7.4 | 3.0 | 3.2 | 6.9 | 3.3 | 7.7 | 5.3 | 48.8 |
|  | *XP* | 0 | 0 | 0 | 0 | 0 | 0 | 0 | 0 | 0 | 100 |
|  | *PXP* | 0 | 0 | 0 | 0 | 0 | 0 | 0 | 0 | 0 | **100** |
| **HC**<br>*fit n=42* | *PP* | 4.5 | 4.5 | 4.5 | 3.8 | 2.8 | 5.3 | 2.8 | 5.5 | 5.8 | 60.5 |
|  | *XP* | 0 | 0 | 0 | 0 | 0 | 0 | 0 | 0 | 0 | 100 |
|  | *PXP* |  |  | 0 | 0 | 0 | 0 | 0 | 0 | 0 | **100** |
| **PSZ**<br>*fit n=31* | *PP* | 11.6 | 8.2 | 11.3 | 4.9 | 7.9 | 9.7 | 8.9 | 10.4 | 4.9 | 22.3 |
|  | *XP* | 5.5 | 1.1 | 4.8 | .1 | 1.0 | 2.5 | 1.7 | 3.3 | .1 | 80.0 |
|  | *PXP* | 10.0 | 10.0 | 10.0 | 10.0 | 10.0 | 10.0 | 10.0 | 10.0 | 10.0 | **10.1** |

### Decision models

Values (RL) or posterior means (HGF) were transformed to choice probabilities by using the softmax (logistic sigmoid) function (see supplementary material). In binary choice tasks with anti-correlated reward-probabilities such as ours, there is choice perseveration independent of learning or inference that differs between win and loss trials. We captured this by estimating parameters for win and loss trials that reflect this difference in choice perseveration (ρ_win_, ρ_loss_). Models which included inverse decision noise β as a free parameter had lower evidence (see supplementary material) than models where this was fixed to unity (β = 1.) We also tested the possibility that volatility is directly linked to choice probabilities by letting third-level trial-by-trial volatility (HGF3) serve as the inverse decision noise (see supplementary material). Because this introduced a volatility scale anchored in observed behavior (switching or staying), it allowed for estimating the mean of the subjective *a priori* belief about initial volatility at the third level, 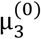, as a parameter of HGF3. This cannot be applied to RL or HGF2 because they do not feature inference on volatility. This led to two additional models (HGF3-SU-V or HGF3-DU-V), resulting in a total of ten models. For model fitting, see supplementary material.

### Model selection

The negative variational free energy (an approximation to the log model evidence) was used for random-effects Bayesian Model Selection (BMS)^45^. The protected exceedance probability (PXP) governed our model selection which protects against the “null” possibility that there are no differences in the likelihood of models across the population^46^. We also examined whether the models explained the data better than chance^17, 47^. A subject was classified as ‘not fit better than chance’ in case the log-likelihood of the data relative to the number of trials did not significantly differ from chance (see supplementary methods). Simulations of the task were run using the inferred parameters to reproduce the observed data.

### Model parameters

Parameters of the winning model were compared between groups using t-tests or the non-parametric Mann-Whitney-U-Test if assumptions of normality were violated (Kolmogorov-Smirnov Test). Bonferroni-correction was applied according to the number of parameters.

### Statistical analysis of fMRI data

Using the general linear model approach in SPM8, an event-related analysis was applied. On the first level, one regressor spanned the entire trial from cue to outcome as in a previous study^38^. We added the following five modeling-based trajectories as parametric modulators (not orthogonalized) to best capture different aspects of the hierarchical inference process: second- and third-level precision-weighted PEs (ε_2_, ε_3_), which were time-locked to the outcome, and precision weights (ψ_2_, ψ_3_) as well as the third level volatility (µ_3_). All regressors spanned the entire trial and changed at outcome accordingly to PE updates identical to ^38^. Regressors were convolved with the canonical hemodynamic response function in SPM8 and its temporal derivative (see supplementary methods). For second-level analysis, a random-effects ANOVA model including contrast images of the five modeling-based trajectories (precision-weighted PEs (ε_2_, ε_3_), precision weights (ψ_2_, ψ_3_) and the third-level volatility (µ_3_) and the factor group was estimated.

## Results

### Behavioral data

Repeated-measures ANOVA on “correct” choices showed that performance differed between phases (dropping in the reversal phase, main effect phase F=23.74, p<0.001). PSZ chose the better card less frequently irrespectively of task phases (Figure 1C, main effect group F=14.52, p<0.001, phase x group F=1.87, p=0.16). The factor phase was dropped from further analyses.

Repeated-measures ANOVA on the probability of choice repetition showed that all participants stayed more with the previous action after rewards compared to losses (main effect feedback F=369.80, p<0.001) and that PSZ switched more, independently of feedback from the previous trial (Figure 1D, main effect group F=27.77, p<0.001, feedback x group F=0.02, p=0.89).

### Computational modeling: model selection

Random-effects BMS^45^ revealed a three-level HGF with double-updating and third-level environmental volatility linked to decision noise (Figure 2) as the most plausible model (HGF3-DU-V, PXP=99.5%, for PXPs of all models, see Table 1). This model (HGF3-DU-V) was superior in HC (PXP=100%, Bayes omnibus risk [BOR] = 0) and remained first-ranking in PSZ (PP=21.4%, XP=74.0%). In PSZ, there was no convincing evidence that models performed differently from each other (BOR=1, all PXP = 10.0%).

15 PSZ and 1 HC were not fit better than chance by any model (Figure 3A). Neither when considering all PSZ (= PSZ-fit + PSZ-nofit) nor PSZ-fit alone did BMS reveal a clearly superior model (both times BOR=1, Table 1). However, the identification of PSZ-fit ensures that individuals included in further modeling-based analyses are fit better than chance by every model (i.e., equally ‘good’ instead of equally ‘poor’ models).

**Figure 3.**
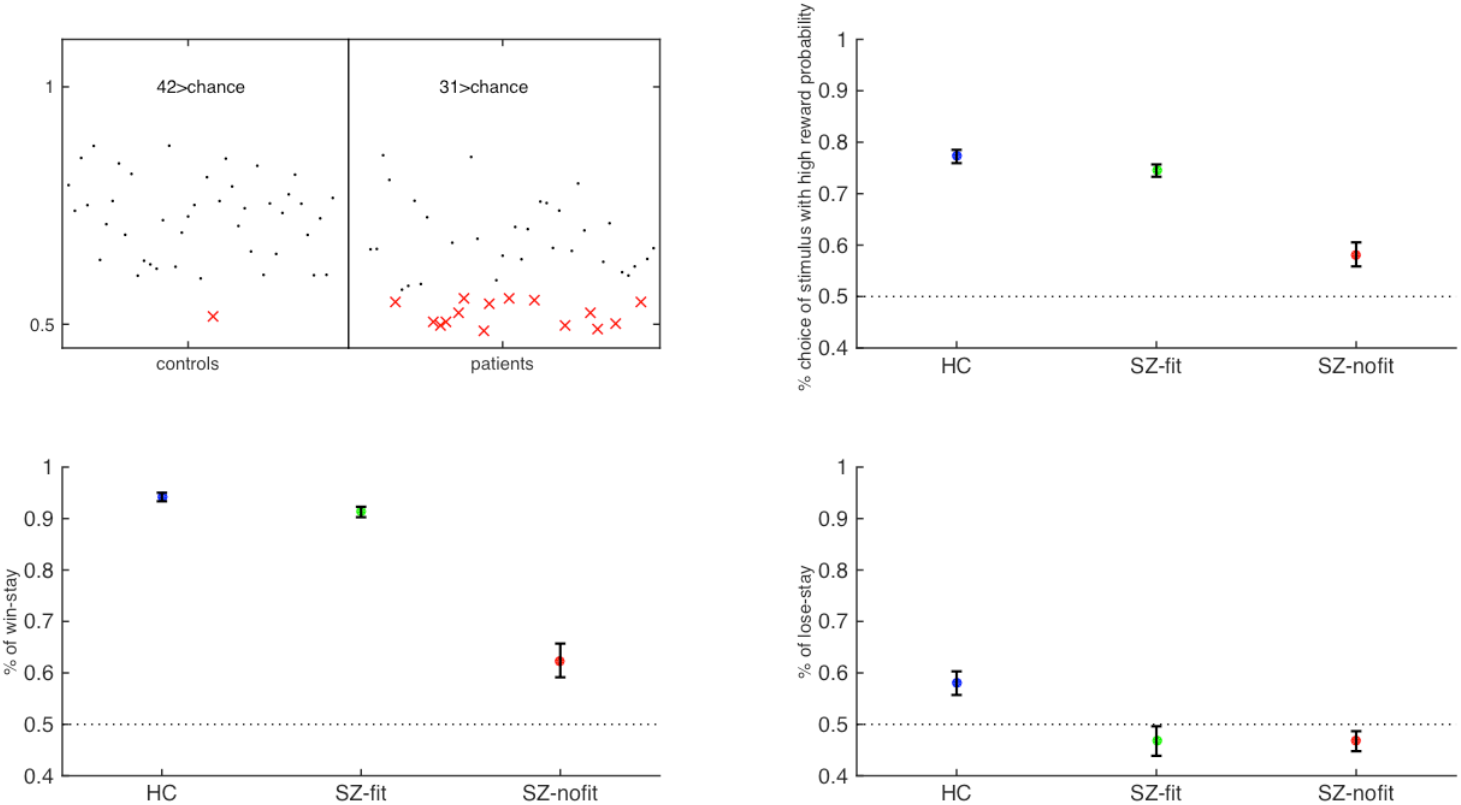
(A) Classification above (black dots) and beIow (red crosses) chance and its influence on overall choice performance (B). There was a main effect of group (F=32.63, p<0.001). PSZ-nofit (red) showed overall poor performance, (panel B, HC vs. PSZ-nofit, t=7.04, p<0.001, PSZ-fit vs. PSZ-nofit, t=6.90, p<0.001), while PSZ-fit (green) performed comparably to HC (blue, Figure 2B, t=1.51, p<0.14). Analysis of win-stay (C) and lose-stay (D) behavior showed a group x feedback interaction (F=20.68, p<0.001). There was a pronounced reduction of win-stay behavior in PSZ-nofit only (C, PSZ-fit vs. PSZ-nofit, t=10.74, p<0.001), while reduced lose-stay behavior characterized both groups of PSZ (D, PSZ-fit vs. PSZ-nofit, t=0.01, p=0.99, (D) This group x feedback interaction was also significant when comparing only HC and PSZ-fit.

### Revisiting behavioral data

Based on this heterogeneity in PSZ regarding absolute model fit, we revisited choice data with respect to three groups (HC, PSZ-fit, PSZ-nofit). There was a main effect of group on “correct” choices (Figure 3B, F=32.63, p<0.001). PSZ-nofit showed performance around chance levels (Figure 3B, HC vs. PSZ-nofit, t=7.04, p<.001, PSZ-fit vs. PSZ-nofit t=6.90, p<0.001), while PSZ-fit had performance comparable to HC (Figure 3B, t=1.51, p=0.14). The analysis of win-stay and lose-stay behavior revealed a group x feedback interaction (F=20.68, p<.001). This resulted from a pronounced reduction of win-stay behavior in PSZ-nofit only (PSZ-fit vs. PSZ-nofit, t=10.74, p<0.001, Figure 3C), while reduced lose-stay was not significantly different between PSZ-fit and PSZ-nofit (t=0.01, p=0.99, Figure 3D). A group x feedback interaction was also significant when comparing only HC and PSZ-fit (F=6.79, p=0.01), next to significant main effects of feedback (F=636.30, p<0.001) and group (F=10.22, p=0.01). This difference between HC and PSZ-fit was driven by switching after loss (Figure 3C & D).

In an exploratory analysis of six cognitive tests, only measures of verbal memory and working memory differed between the two groups of PSZ, i.e. were more impaired in PSZ-nofit compared to PSZ-fit (see supplementary results). This suggests that PSZ-fit and PSZ-nofit mapped on distinct cognitive profiles. Because poor fit hinders the interpretation of modeling-based behavioral and neuroimaging analyses in the PSZ-nofit subgroup, all subsequent modeling-based results are reported based on HC (n=42) and PSZ-fit (n=31) only.

### Computational modeling: parameters

Comparison of parameters of HGF3-DU-V (Table 2, Figure 4) revealed that the estimated mean of the *a priori* belief about initial environmental volatility 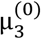 was higher in PSZ (z=3.15, p<0.01, Figure 4A). Trial-by-trial environmental volatility was more strongly coupled with lower-level updating, as demonstrated by higher κ in PSZ (z=2.51, p<0.01, Figure 4B). The evolution rate of environmental volatility *ω*_3_ did not differ significantly between groups (z=0.73, p=0.47). To illustrate the effects of differences in parameters on behavior after losses (when PSZ-fit showed increased switching), we analyzed the trajectory of *µ_3_*in a mixed-effects regression model with group and feedback as predictors. This revealed higher *µ_3_*in PSZ-fit overall. Across groups, *µ_3_* was higher after losses compared to rewards, which was more pronounced in PSZ (resulting from enhanced coupling between higher and lower levels κ). For statistics, see supplementary results (S-Figure 2).

**Figure 4.**
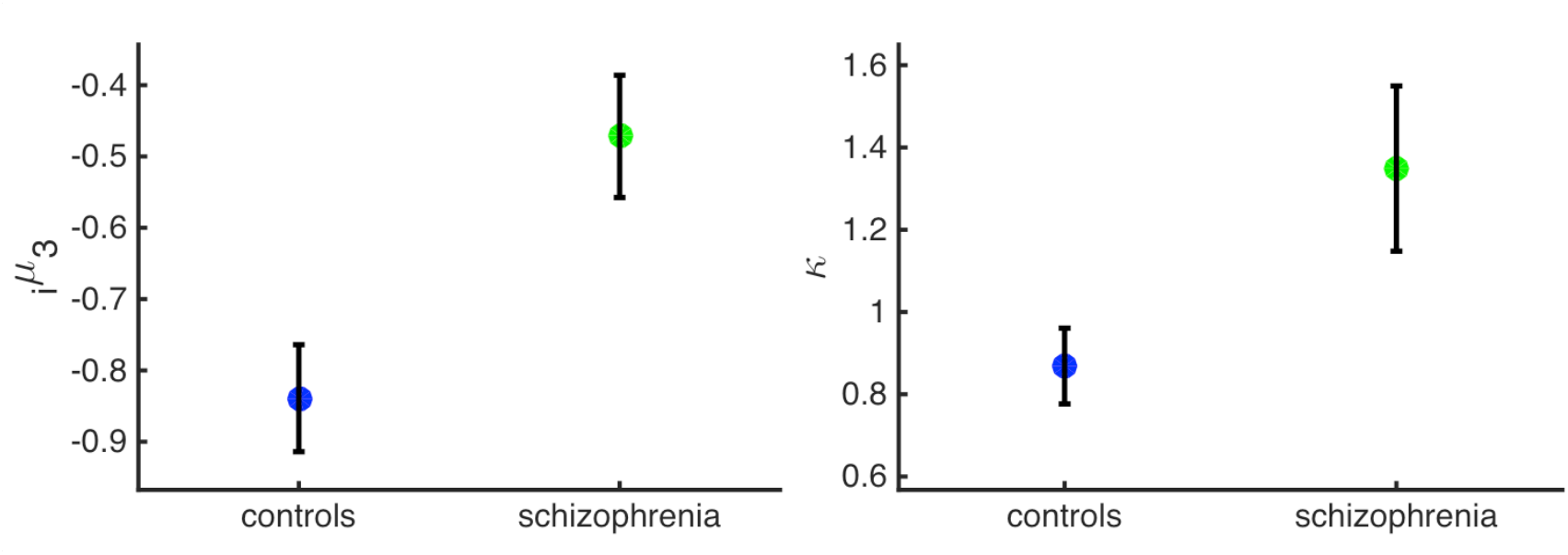
(A) The initial estimate of environmental volatility is significantly higher in PSZ-fit (n=31) as compared to HC. (B) The coupling between the third level (environmental volatility) and the second level is significantly stronger in PSZ-fit (n=31) compared to HC.

**Table 2.** Between-group comparisons of model parameters using t-tests or the non-parametric Mann-Whitney-U-Test if assumptions of normality were violated. Bonferroni-correction was applied according to the number of parameters (5, p<0.01). HC = healthy controls, PSZ = patients with schizophrenia.

|  | HC (n=42) | PSZ (n=31) | Test statistic |
| --- | --- | --- | --- |
| *Learning model* |  |  |  |
| $\mu_3$ | $-0.84 \pm 0.49$ | $-0.47 \pm 0.48$ | $z=3.15, p<.01$ |
| $\kappa$ | $0.87 \pm 0.60$ | $1.35 \pm 1.12$ | $z=2.51, p<.01$ |
| $\omega_3$ | $-6.00 \pm 0.02$ | $-5.99 \pm 0.05$ | $z=0.73, p=0.47$ |
| *Decision model* |  |  |  |
| $\rho_{\text{win}}$ | $0.97 \pm 0.57$ | $1.08 \pm 0.49$ | $z=0.88, p=0.38$ |
| $\rho_{\text{loss}}$ | $0.08 \pm 0.31$ | $-0.12 \pm 0.44$ | $z=2.37, p=0.02$ |
| *Replication sample: Schlagenhauf et al. 2014* |  |  |  |
|  | HC (n=24) | non-medicated PSZ (n=24) |  |
| *Learning model* |  |  |  |
| $\mu_3$ | $-1.17 \pm .61$ | $-.43 \pm .70$ | $z=3.88, p<.01$ |
| $\kappa$ | $.61 \pm .58$ | $1.56 \pm 1.21$ | $z=3.46, p<.01$ |
| $\omega_3$ | $-6.09 \pm .02$ | $-5.99 \pm .06$ | $z=1.35, p=.18$ |
| *Decision model* |  |  |  |
| $\rho_{\text{win}}$ | $.70 \pm .74$ | $.43 \pm .66$ | $z=1.35, p=.18$ |
| $\rho_{\text{loss}}$ | $-.07 \pm 0.59$ | $-.28 \pm .77$ | $z=1.05, p=.30$ |

### Computational modeling: reproducing observed behavior

Simulating data based on the inferred parameters of HGF3-DU-V (42 HC, 31 PSZ-fit, 10 simulations per subject) showed that PSZ-fit switched more than HC (main effect of group, F=11.17, p<0.001) and showed a pronounced tendency to switch after losses (group x feedback F=7.68, p=0.01). Between-group findings on behavioral data were fully reproduced, which yields an important validation of the model’s ability to capture the observed data.

### Computational modeling: replication in non-medicated PSZ

We tested our model (HGF3-DU-V) in an independent sample of non-medicated PSZ (n=24) and HC (n=24), who performed another reversal-learning task^17^. This replicated between-group findings and remained significant when excluding participants not fit better than chance (23 HC, 13 PSZ). For statistics, see Table 2.

### Relation to symptoms

We explored the relation between the two parameters that differed between groups with different measures of cognition (n=6) and clinical measures (n=7) within PSZ (S-Table 1) applying Bonferroni-correction (p<0.0083). In PSZ, a higher initial belief about volatility 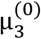 was associated with reduced executive functioning and cognitive speed (TMT B: r=-0.56, p=<0.001; DSST: r=-0.56, p<0.001, S-Table 3, S-Figure 3). These correlations were not present in the healthy control group. For all explorative correlations, see supplementary material (S-Table 3).

### FMRI – task effects (pooled across groups)

Activity related to ε_2_ (a precision-weighted RPE) peaked in bilateral ventral striatum and ventromedial PFC among other regions (p-FWE_wholebrain_<0.05, Figure 5A, S-Table 4) including the midbrain (p-FWE_midbrain-voi_<0.05, S-Table 9), a well-known network associated with RPEs. In contrast, third-level precision-weighted PE (ε_3_) was associated with activity in prefrontal, parietal and left insular regions (Figure 5A, S-Table 5). Environmental volatility (µ_3_) co-varied with activation in bilateral insula, cingulate cortex, parietal cortex, middle temporal gyrus, globus pallidus, thalamus as well as superior, middle and inferior frontal gyrus (Figure 6A, S-Figure 5, S-Table 8). For more details on group-level fMRI effects including activity specific for each group outside the effect of each regressor combined for HC and PSZ, please see supplementary results.

**Figure 5.**
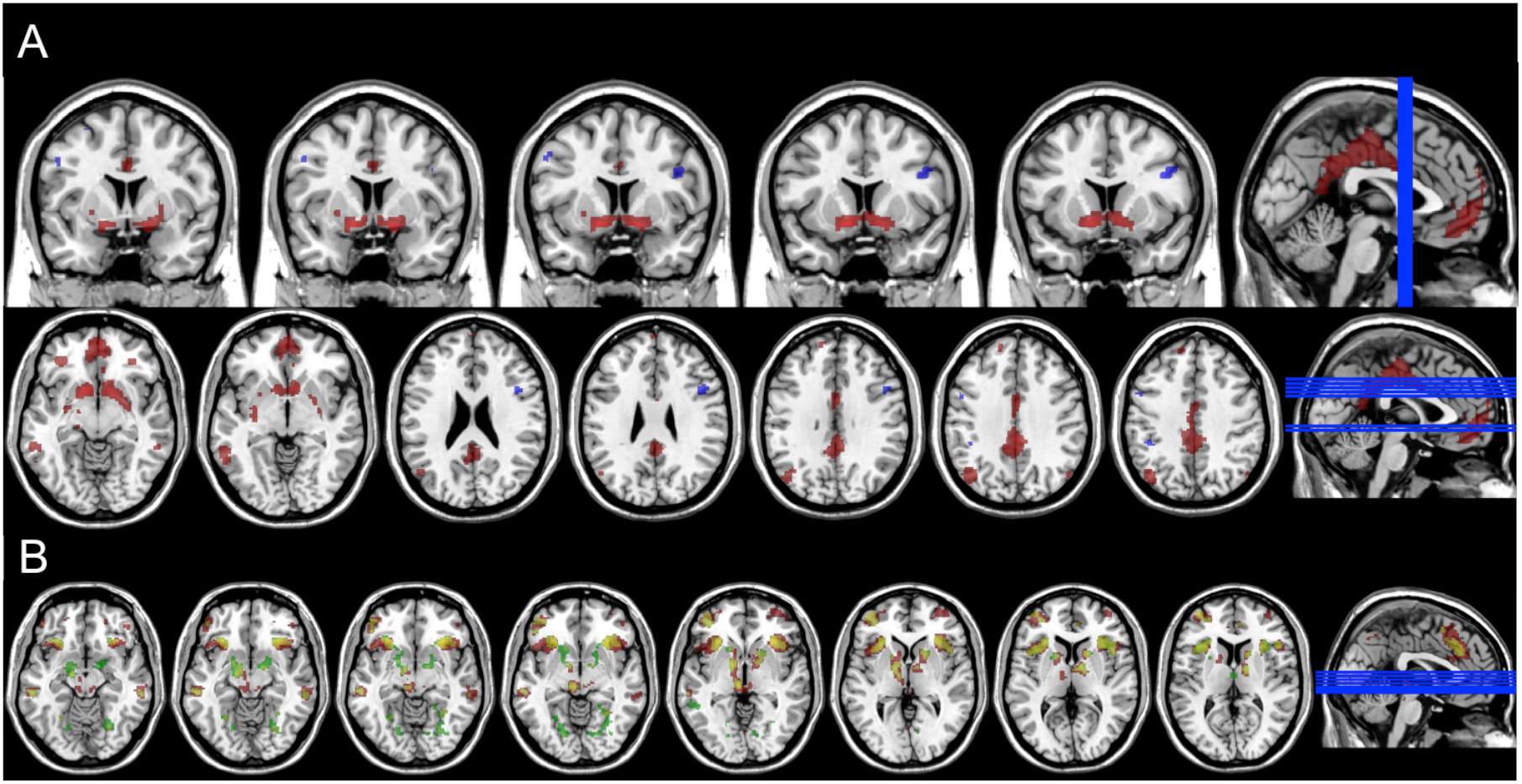
BOLD signal across all participants related to (A) precision-weighted prediction errors from the second (red) and third (blue) level and (B) to precision weight from the second (red) and third (green) level, overlap in yellow. Both at p-FWE_wholebrain_<0.05, k=10.

**Figure 6.**
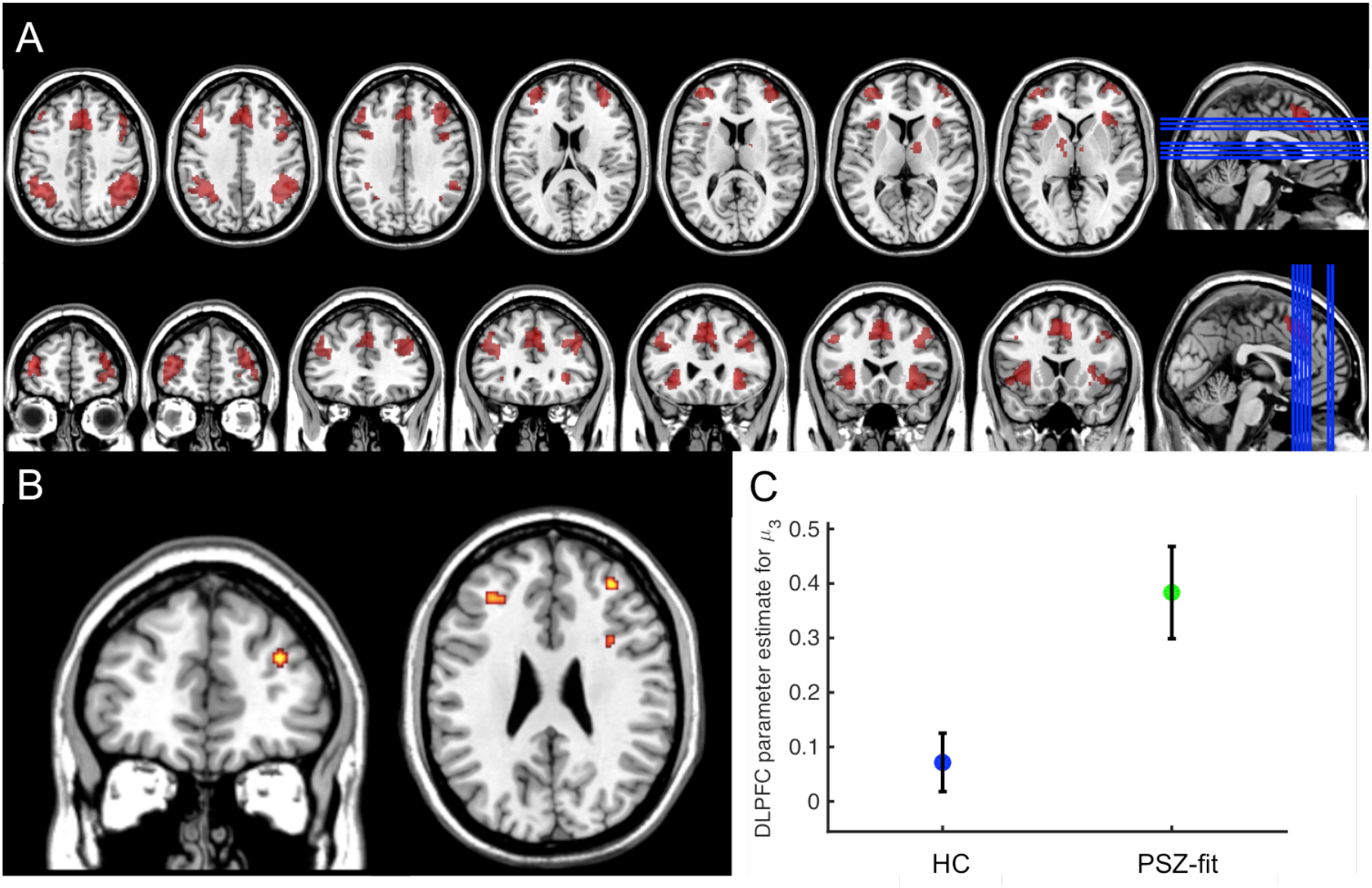
(A) Across all participants, BOLD signal related to volatility estimate from the third level at p-FWE<0.05 for the whole brain, k=10. (B) µ_3_-related BOLD signal in the dorsolateral prefrontal cortex (dlPFC) differs between PSZ-fit (n=31) and HC (F-contrast displayed at p<.001 uncorrected; corrected for main effect of µ_3_ over all participants, [x=34, y=44, z=24], F=19.89, z=4.24, p=0.038). (C) A post-hoc analysis of regression parameter estimates at the peak of the group difference [x=34, y=44, z=24] showed that this was driven by heightened activation related to environmental volatility in dlPFC of PSZ-fit (n=31) compared to HC (t=4.46, z=4.4, p=0.019).

### FMRI – between-group effects

We conducted *between-group comparison of the covariance between the modelling regressors derived from the best-fitting model and BOLD* response within SPM. For the regressor of environmental volatility *µ_3_*, a group difference between HC and PSZ was found in right dlPFC (F-contrast, using a mask representing the average effect of *µ_3_* over all participants for correction of multiple comparison, [x=34, y=44, z=24], F=19.89, z=4.24, p_FWE_=0.04, Figure 6B). Post-hoc analysis revealed stronger activity related to volatility in dlPFC of PSZ compared to HC (t=4.46, z=4.4, p_FWE_=0.02, Figure 6C). There was no significant difference between groups for any other regressor.

## Discussion

To the best of our knowledge, this is the first study to apply hierarchical Bayesian learning to choice and fMRI data of PSZ during reward-based decision-making. We present two main findings: Firstly, our modeling suggests that medicated PSZ acted under an *a priori* enhanced higher-level belief about initial environmental volatility and increased coupling between higher and lower levels of learning, which leads to enhanced lower-level belief updating about action-outcome contingencies. This provides a computational account of choice switching as previously observed in PSZ^8, 12–17^. Using parameters of the winning model to simulate new data, we fully reproduced observed patterns in the behavioral data, and, we replicated our findings on parameters in an independent cohort of non-medicated PSZ. Secondly, medicated PSZ displayed higher dlPFC activity related to environmental volatility, which points towards a prominent role of this region in promoting instable behavior in PSZ.

As reported previously^8, 12–17^, PSZ showed enhanced choice switching and we demonstrate a possible underlying mechanism: an enhanced initial belief about the environmental volatility and a stronger coupling of volatility and lower-level learning of action-outcome contingencies. This has two consequences. Firstly, PSZ had an overall stronger tendency to switch (enhanced initial belief about volatility). Secondly, lower-level beliefs fluctuated more strongly and led to increased choice switching particularly after (irrelevant) losses. Thus, PSZ inferred more contingency changes in this dynamic task environment, which are putatively signaled through losses. Enhanced estimates of changes in context probabilities wer also found in PSZ in the non-reward domain^48^. In contrast to our finding of enhanced initial belief about volatility, Powers et al. (2017) probed conditioned hallucinations and reported stronger lower-level priors about perceptual inputs combined with reduced evolution of volatility in hallucinating participants. A possible explanation may be that alterations of volatility estimates differ with regard to the investigated functional domain, potentially related to different symptom dimensions. As suggested by our finding, cognitive beliefs about the structure of the environment appear to be more unstable, while perceptual beliefs about sensory inputs appear to be overly stable and not appropriately adjusted following changes in the environment^31^.

Because our model revealed consistent results in medicated und non-medicated PSZ, elevated beliefs about environmental volatility may represent an important mechanism of impaired flexible decision-making. In a similar vein, an inability to stabilize behavior according to an internal model of action-outcome contingencies was found after administration of ketamine in healthy controls^49^, which is in line with the assumption that reduced (prefrontal) NMDA receptor functioning^27, 50^, may lead to aberrant cortical information processing^51^. In line with this idea, we found a stronger association in PSZ than in HC of beliefs about volatility with BOLD activity in dlPFC. However, in our fMRI study we cannot not infer about involved neurochemical systems. On the behavioral level, beliefs about higher volatility, in our model directly linked to decision noise, can lead to more stochastic behavior (in our task overall more choice switching). We therefore suggest, that heightened neural representation of volatility may generate more stochastic behavior, although in our correlational and cross-sectional study we cannot ascertain a causal link. We found a negative correlation of the initial belief about volatility with independent neuropsychological measures of executive functioning and cognitive speed in PSZ, thereby emphasizing its dysfunctional character, while these association were not observed in the healthy control group.

PSZ’s heightened belief about volatility may render a system (hyper-)sensitive to any new input^51, 52^, thereby impeding the detection of regularities in probabilistic environments and leading to (hyper-)flexible updating in response to (irrelevant) information. Meta-analyses showed reduced prefrontal activity in PSZ compared to HC for (‘relevant’) task vs. (‘irrelevant’) conditions across multiple cognitive measures^53, 54^. On the one hand, prefrontal dysfunction in PSZ may contribute to an enhanced higher-level belief about volatility (e.g., by impairing the detection of higher-level regularities) and such beliefs about volatility might be assigned with enhanced precision (potentially in a compensatory manner). On the other hand, lower-level beliefs may be more unstable and presumably assigned with lower precision leading to distinct aberrant experiences and behaviors depending on the tested domain with so far most evidence coming from perceptual processing^55^.

Aberrant cortical processing was theorized, at least in non-medicated PSZ, to increase striatal dopamine turnover^27, 50^, which might interfere with striatal and midbrain RPE signals. Indeed, in non-medicated PSZ striatal RPE activity was found to be reduced^17, 56^. In RL accounts^24^, enhanced spontaneous phasic dopamine transients could highlight irrelevant stimuli and disturb the signaling of (relevant) RPEs. In our medicated sample of PSZ, no significant differences in striatal activations to RPEs were observed, which is in line with reports of absent differences in striatal RPEs in medicated PSZ^14, 15^. This suggests medication status as an important factor relating to striatal RPE signaling similar to medication effects on striatal reward anticipation in PSZ^57–59^. In hierarchical Bayesian learning, representation of lower-level PEs may be similar in patients and controls but potentially reduced precision of lower-level beliefs might highlight irrelevant inputs (e.g. resulting in choice switching). This could, at least in theory, result from a common aberrant prefrontal process, as discussed above, but also from an effect of antipsychotic D2 receptor antagonists in the striatum^29^. However, we did not observe group differences in midbrain and striatum for precision weights and precision-weighted PEs. While our data indicate a disrupted higher-level process with evidence from behavioral modeling and fMRI, disturbed lower level processes are supported by our behavioral modeling but not by the presented fMRI data.

### Limitations

Firstly, the lack of clear superiority of any model for the data from PSZ, as well as the substantial number of PSZ in which no model fitted better than chance needs to be considered. Excluding these patients from further modeling-based analyses can be considered restrictive and may reduce the generalization of results. However, two subgroups were identified with (task-independent) different cognitive profiles which contributes to disentangling heterogeneity across schizophrenia patients -- a fundamental challenge for psychiatric research^60^. By controlling whether subjects’ performance can actually be interpreted as assumed by the theories, we eliminated a potential key confound. Nevertheless, it remains problematic if clinical groups differ in how well they are described by models of interest, as observed in our study because parameters are conditional on the model. This impedes the interpretation of differences in parameters across groups (see ^61^ for a discussion of this problem). In principle, Bayesian model averaging^62^ can help, but this is not established for non-nested models as used here. Future studies might implement tasks with adaptive difficulty to reduce the number of patients whose behavior cannot be explained by any model potentially due to excessive cognitive demands. Furthermore, the relation between different Bayesian modeling approaches such as the HGF and active inference models should be explored^63^.

Secondly, we suggest that overestimating volatility is one possible explanation for choice switching. However, in our model, volatility partly determines decision noise. This limits the differentiation between the concepts of volatility and exploration and limits interpretability of volatility estimates to some extent. The finding that this model performs best is an indication that our data do not fully support a strong distinction between volatility and exploration. Additional task-based manipulations would be required to overcome this^37^, which will most likely involve longer tasks than our patient-friendly fMRI task (<15min, 160 trials). The formulation of our response model suggests that results are still informative. We control for overall differences in stickiness with two parameters that change the shape of the decision function differently for wins and losses and exert a bias towards repeating or switching responses, irrespectively of learned expectations (see supplementary material). Therefore, decision noise is not only determined by trial-wise volatility but also by subject-specific traits that are expressed in a condition-specific fashion. This disentangles volatility and decision noise to some degree.

Thirdly, this is a case-control study which fundamentally limits the inferences that can be drawn from the results, e.g. the development of inappropriately high initial beliefs about volatility over the course of illness, its stability over time and its relation to broader cognitive deficits consistently found in PSZ.

In summary, we present a computational mechanism putatively underlying instable behavior in PSZ: a stronger coupling of heightened beliefs about environmental volatility with lower-level learning, which was present in medicated und non-medicated PSZ. In medicated PSZ, this was accompanied by enhanced activity related to environmental volatility in dlPFC. Future studies should aim to test specificity of the presented results for PSZ and overcome the limitation of the lack of longitudinal clinical data. Computational modeling may aid in the identification of subgroups of PSZ^64^ and potentially inform the prediction of treatment response to antipsychotic drugs by aiming to dissect the important biological heterogeneity and inter-individual differences among patients^65^. These steps towards clinically useful procedures will require carefully designed prospective studies in the framework of Computational Psychiatry^29, 66–68^.

## Author contributions

LD, CM, KES & FS designed the study. LD, RB, TK & JK performed research. LD & RB analyzed the data, CM & FS supervised data analysis. LD & RB wrote the initial version of the manuscript. LD, RB, TK, JK, CM, KES, AH, FS read and corrected versions of the manuscript.

## Acknowledgements

This study was supported by the Max Planck Society and grants from the German Research Foundation awarded to FS (DFG SCHL1969/1-2, DFG SCHL 1969/3-1, SCHL1969/4-1). KES acknowledges support by the René and Susanne Braginsky Foundation and the University of Zurich. Data from this study was presented at the following conferences: 71^st^ annual meeting of the Society for Biological Psychiatry, Atlanta, USA, May 12-14, 2016; The 6th Biennial Schizophrenia International Research Society Conference (SIRS), Florence, Italy, 4-8 April 2018; 11^th^ Forum of Neuroscience, Berlin, Germany, 7-11 July 2018.

## Supplemental Methods

### Participants and instruments

Handedness was assessed with the Edinburgh Handedness Inventory. Cognitive functioning was examined with six subscales selected from the Wechsler Adult Intelligence Scale^1^. Symptoms were assessed with the Positive and Negative Syndrome Scale (PANSS) and with delusion and anhedonia items from the Scale for the Assessment of Positive Symptoms and Negative Symptoms (SAPS/SANS). Dosage of antipsychotics was standardized by chlorpromazine equivalents^2^.

### Task

Participants performed a task requiring flexible decision-making^3–5^ (Figure 1A). In 160 trials, individuals decided between two cards each showing a different visual stimulus. The right or left location of stimuli was randomized over trials. After left or right button press (max. 1.5s), the selected card was highlighted and a monetary win (10 Eurocent coin) or a monetary loss (crossed 10 Eurocent coin) was shown for .5s. In the inter-trial interval (exponentially distributed jitter, range 1s-12.5s), a fixation cross was presented. If no response occurred in time, the message “too slow” appeared. One of two cards was initially assigned with a reward probability of 80% and a loss probability of 20% and vice versa for the other card. The task had a simple higher-order structure (Figure 1B): an anti-correlation between the reward probabilities; whenever one card was associated with a probability of 80%, the other card would be associated with a probability of 20%. Reward contingencies were stable for the first 55 trials (‘pre-reversal’) and for the last 35 trials (‘post-reversal’). During the ‘reversal’ phase, contingencies changed four times, after 15 or 20 trials, respectively.

Before MRI, participants were informed that one card had a superior chance of winning money. They were told that depending on their choice they could either win or lose 0.10€ per trial and to win as much as possible as the total gain was paid out. 20 training trials were performed with different cards and without reversal. After training, participants were instructed that reward probabilities could change and to track such changes to win as much as possible. No other information on reversals or the anti-correlated task structure was given.

### Computational models of learning

Models of learning comprised RL and the HGF. In all models, RPE δ is used to update expectations for the chosen stimulus:

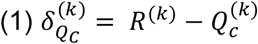

Using the notation of RL, 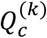 represents the expectation for receiving reward or loss by choosing the card *c* in trial κ. R denotes the received outcome. *RPE δ* reflects the difference between received rewards and expectations. The update of 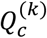 is equal to the RPE 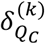 weighted by the learning rate α:

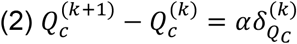

Learning rate α ∈ [0, 1] reflects a constant weight on RPE determining the change in values throughout trials κ. With learning rates close to 1, prediction errors (PEs) strongly affect expectations, while learning rates close to 0 lead to little influence of PEs on expectations.

In the generative model underlying the HGF, the evolution of states x_i_ (with i indexing the level) is defined as a Gaussian random walk (except for *i* = 1, where *x*_1_ represents the binary outcome of the trial: reward or loss). The change in each *x*_i_ can be inferred using the HGF’s update equations. These provide trial-wise inference on states *x_i_*, represented by the posterior mean µ_i_ and variance σ_i_ at each level. It is important in this context to distinguish between the underlying generative model (containing states *x_i_*) and the HGF properly (i.e, the inference process containing state representations µ_i_ and σ_i_). In our implementation, action-outcome contingencies *x_2_* evolved as a Gaussian random walk; the probability for a choice being rewarded (i.e., *x_1_* = 1) at a given trial κ is 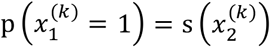, where s(·) is the logistic sigmoid function. The step size of the Gaussian random walk of *x_2_* depends on the next higher level *x_3_*:

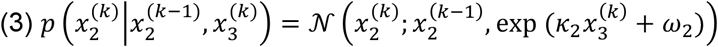

The variance of this conditional distribution is parameterized by *ω_2_* and κ. The term 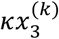 couples the third-level environmental volatility 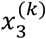 with the second level. 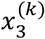 evolves in the same manner, except that the variance of the Gaussian is a constant *ω_3_* (because there is no higher level):

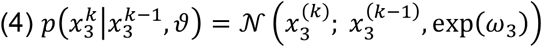

In our implementation, *ω_2_* was fixed at the second level because we were particularly interested in inferring the parameters that influenced subject-specific estimates of environmental volatility at the third level (*ω_3_*) and its coupling with the lower level (κ). Variational inversion of this generative model results in the HGF. This inversion shows that trial-by-trial updates of posterior means at each level *i* are proportional to the PE from the level below weighted by a precision ratio (compare equation 2 for the RL equivalent), where precision 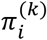 is defined as inverse variance 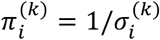.

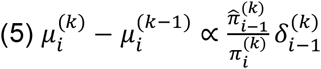

For an exact derivation of precision weights and precision-weighted PEs, we refer to previous methodological papers^6, 7^. In addition to the three-level HGF (HGF3), we also included a simpler two-level variant (HGF2) to test whether the full representation of volatility in the HGF3 was superior in explaining behavior.

In these models, HGF and RL update expectations of the chosen card only (“single-update”, SU) and the expectation about the unchosen card remains unchanged.

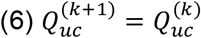

Correspondingly, for the HGF:

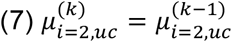

Based on the anti-correlated task structure, we implemented a variant of each learning model updating values (RL, equation 8) or posterior means (HGF, equation 9) of the unchosen card simultaneously (“double-update”, DU), which can be written as:

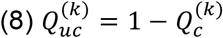

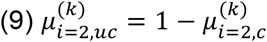

### Decision models

For each trial κ, Q^(k)^ (RL) or 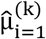 (HGF) were transformed to choice probabilities by using the softmax model:

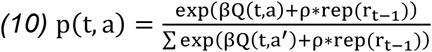

In binary choice tasks with correlated reward-probabilities, there is strong autocorrelation of choices (perseveration). In our decision model (equation 10), this is captured by a parameter ρ. This parameter changes the inflection point of the sigmoid function, biasing choice predictions towards an overall tendency to stay or switch irrespectively of the learned expectations. We used separate parameters ρ_win_ and ρ_loss_ to reflect differences in choice perseveration after rewards and losses separately. This is in line with previous observations that choice perseveration differs strongly after rewards and losses. Note that this implementation of capturing choice perseveration is equivalent to fitting parameters for rewards and losses as part of the learning model (cf. previous work^8, 9^), which changes the asymptote of expectations in the learning model instead of the inflection point of the sigmoid decision model. We name these decision models ‘REP’. In each of the ten models reported in the main manuscript the parameter β was fixed to 1 to avoid overparameterization. In addition, we also estimate all learning models with a decision model with no such repetition parameters but β as a free parameter (‘BETA’). The parameter β captures the steepness of the sigmoid and thereby reflects (inverse) decision noise by determining how tightly choice probabilities follow learned reward expectations. The ten learning models with β capturing inverse decision noise provided an overall inferior account of the data when performing a family comparison (REP XP_all_=.8550, BETA XP_all_=.1450; REP XP_HC_=.8579, BETA XP_HC_=.1421; REP XP_SZ_=.7316 BETA XP_SZ_ = .2684, [XP: exceedance probability]).

We also tested the possibility that the parameter β was directly influenced by trial-by-trial environmental volatility from the third-level of HGF3 by introducing a trial-wise 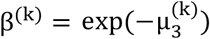. Because this equates the inverse decision temperature to the inverse of the exponentiated log-volatility, it amounts to letting the volatility serve as the decision temperature.

### Model fitting

HGF toolbox 3.15 http://www.translationalneuromodeling.org/tapas/; ^40, 41^ with a quasi-Newton optimization algorithm was used for model fitting. For prior means and variances of parameters, see S-Table 2.

### Model selection

Using random-effects Bayesian Model Selection (BMS)^10^, which accounts for heterogeneity across subjects by treating the model as a random variable in the population, a posterior model probability (PP) is provided for each model, i.e., the probability that the data from a randomly chosen subject are best explained by this model. The certainty about this probability is quantified by the exceedance probability (XP), i.e. the probability that this model is more likely than any other model considered. One can protect against the “null” possibility that there are no differences in the likelihood of models across the population^11^; this yields the protected exceedance probability (PXP), which was the metric that governed our model selection.

### Classification of subjects not fit better than chance

In choice tasks like ours, a subject can be classified as fit better than chance when the geometric mean likelihood per trial (given by exp(log-likelihood/ntrials)) significantly exceeded 0.5, which corresponds to p<0.05 when performing a binomial test. This procedure was applied to all individuals to avoid the possibility that between-group differences in model parameters are confounded by differences in model fit^4, 8, 12–14^. An alternative would be to capture (undirected) noise with a parameter in the decision model, as suggested previously^15^. While this is an elegant way to formulate the problem, it is less straightforward to identify subjects whose behavior cannot be sufficiently explained by non-random mechanisms (in the models considered). Here, we remove such individuals from statistical comparisons and treat them separately in the analyses of choice data.

### Functional Magnetic Resonance Imaging

Functional Magnetic Resonance Imaging (fMRI) imaging was performed using a 3 Tesla Siemens Trio scanner to acquire gradient echo T2*-weighted echo-planar images with blood oxygenation level dependent contrast. Covering the whole brain, 36 slices were acquired in oblique orientation at 20° to AC-PC line in ascending order with 2.5-mm thickness, 3×3mm² in-plane voxel resolution, 0.5-mm gap between slices, TR=2s, TE=22ms and a flip angle α=90°. Prior to functional scanning, a field map was collected to account for individual homogeneity differences of the magnetic field. T1-weighted structural images were also acquired (TR=1300ms, TE=3.46ms, flip=10°, matrix=240×256, voxel size: 1×1×1mm, slices=170).

### Analysis of fMRI data

FMRI data were analyzed using SPM8 (http://www.fil.ion.ucl.ac.uk/spm/software/spm8/). For preprocessing, images were corrected for delay of slice time acquisition. Voxel-displacement maps were estimated based on field maps. All images were realigned to correct for motion and were also corrected for distortion and the interaction of distortion and motion. The images were spatially normalized into the Montreal Neurological Institute (MNI) space using the normalization parameters generated during the segmentation of each subject’s anatomical T1 scan; spatial smoothing was applied with an isotropic Gaussian kernel of 6mm full width at half maximum. Prior to first-level statistical analysis, data were high-pass filtered with a cutoff of 128s. In the first level model, missed choices were modeled separately. The six movement parameters were included in the model as regressors of no interest as well as the first derivative of translational movement with respect to time. An additional regressor was included censoring scan-to-scan movement >1 mm.

## Supplemental Results

### Clinical distinction between subgroups

When further exploring the subgroups, SZ-nofit did not significantly differ from SZ-fit in any of the seven measures of positive and negative symptoms (all p’s>.3910). Interestingly, SZ-nofit showed significantly lower performance when compared to SZ-fit in terms of certain domains of cognitive functioning such as verbal memory (word list p=.0329), working memory (digit span p=.0271) but not verbal IQ (p=.1024), attention (TMT A p=.5158) or cognitive speed (DSST p=.3091) but trend-wise with respect to executive functioning (TMT B p=.0697). However, this analysis should be considered exploratory as none of the effects reached correction for multiple comparisons for the 6 cognitive tests (p<.0083).

### Computational modeling: parameters

In the decision model, choice repetition after rewards did not significantly differ between groups (z=0.88, p=0.38) while staying after losses was significantly lower in schizophrenia (z=2.37, p=0.02) but did not survive Bonferroni-correction for the model’s five parameters. To illustrate effects of parameters that differed between groups on the third-level environmental volatility, mixed-effects regression were estimated in R. The trial-wise posterior mean 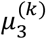 of inferred environmental volatility per subject was the dependent variable while group and feedback were included as independent variables. This model showed a significant main effect of group (t=3.22, p=0.01) due to higher estimates of volatility in PSZ and a main effect of feedback (t=16.124, p<.001) due to higher estimates of volatility after losses compared to rewards. Of most interest, there was also a group x feedback interaction (t=2.50, 0.02). This was driven by a larger difference in environmental volatility after having received losses versus rewards in PSZ as compared to HC (S-Figure 2).

### Computational modeling: recovery of parameters

In addition to showing that data generated from the inferred parameters reproduces the behavioural results (see main manuscript), an additional step is to recover parameters by refitting the generated data. After having done so, we tested parameters, as inferred from generated data, between groups and found the same between-group differences as when inferring the parameters from the observed data (*μ_3_: HC* −1.46 ± 0.60, *PSZ* −1.04 ± 0.55, z=3.09, p<.01; *κ: HC* 0.61 ± 0.30, *PSZ* 0.73 ± 1.12, z=2.15, p=.03; *υ: HC* −6.00 ± 0.02, *PSZ* −6.00 ± 0.02, z=1.40, p=0.16; *ρ*_win_: *HC* 1.04 ± 0.60, *PSZ* 1.28 ± 0.62, z=1.68, p=0.10; *ρ*_loss_: *HC* 0.19 ± 0.27, *PSZ* 0.01 ± 0.38, z=2.40, p=0.02). We also correlated parameters inferred from observed data with parameters inferred from generated data. This resulted in the following correlations coefficient: rρ_win_ =.73, rρ_loss_ =.78, rμ_3_ =.87, rκ =.56, rυ =-03.

### Relation to symptoms

When exploring the relation of the two parameters that differed between groups with measures of cognition (n=6) and psychopathology (n=7) within PSZ, we found were negative correlations between the prior belief about volatility 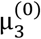 and measures of executive functioning and cognitive speed surviving Bonferroni-correction (p<0.0083, TMT B: r=-0.56, p=<0.001; DSST: r=-0.56, p<0.001, S-Table 3, S-Figure 3). Correlations did not reach significance between cognition and κ nor between the two parameters and any of the seven symptom scales. The strongest, albeit non-significant, correlations were observed between κ and delusions (r=0.30, p =0.10) and between 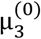 and anhedonia (r=0.32, p =0.09). For all explorative correlations, see supplementary material (S-Table 3).

### FMRI – task effects pooled across groups

In conjunction analysis, ε_2_ and RPE from RL largely overlapped (S-Table 9; S-Figure 4). Significant BOLD responses related to second-level precision weight (ψ_2_) were found in bilateral insula, dorsolateral prefrontal cortex (dlPFC), cingulate cortex, parietal cortex, caudate and thalamus (Figure 4B, S-Table 6). Interestingly, activation associated with third-level precision weight (ψ_3_) estimates was largely (but not entirely) overlapping with that associated with second-level precision weight ψ_2_ (Figure 4B, S-Table 7).

### Task effects unique for either HC or PSZ

In order to identify neural correlates engaged in the task that are unique to PSZ or HC, we report areas that are significantly activated for either HC or PSZ and which are outside the effect of each regressor (both threshholded at p<0.05, FWE-corrected on the whole brain level, clustersize k = 10). For the PSZ group, we found a unique activation for ε2 in the left motor cortex (S-Table 4), for Ψ3 bilaterally in the cerebellum (S-Table 7) and for µ3 in the right dlPFC (S-Table 8). For the HC group, we found a unique activation for µ3 in the medial PFC and in the left inferior frontal gyrus (S-Table 8).

### Neural correlates of reward prediction errors based on Reinforcement Learning

As in previous studies^8^, we also explored correlates of prediction errors using a regressor of reward prediction errors from the RL model. As previously, we found that BOLD signal covaried with RPEs in the ventral and dorsal striatum, posterior cingulate cortex, dorsolateral and lateral PFC, parietal cortex, hippocampus, amygdala and the cerebellum (S-Table 9, S-Figure 3A). There was no significant difference between the groups. However, this signal largely overlapped with the precision-weighted PE at the second level of the HGF (Figure 3A, S-Table 4, S-Figure 3B), which represents, in the case of our tasks, a precision-weighted RPE.

---

**S-Table 1.** Sample Characteristics. Group means with standard deviations and range in brackets are reported; PSZ=Patients with schizophrenia, HC=healthy controls, CPZ=chlorpromazine equivalents; for group comparisons two-sample t-test or χ^2^ tests were used. *=Bonferroni-corrected

|  | PSZ<br>(N=46) | HC<br>(N=43) | Sig. |
| --- | --- | --- | --- |
| Age (years) (46/43) | 35.07 ± 7.596 (22-52) | 34.40 ± 8.24 (22-51) | .69 |
| Gender (46/43) | 14 female / 32 male | 13 female / 30 male | .98 |
| Handedness (44/40) | 38r 4l 2b | 32r 2l 6b | .22 |
| EHI (44/40) | 70.71 ± 53.13 (-100-100) | 71.68 ± 47.55 (-100-100) | .93 |
| Smoking (44/39) | 35 smoking, 9 non-smoking | 15 smoking, 24 non-smoking | <.01 |
| *Cognitive measures* |  |  |  |
| Verbal Intelligence (45/42) | 98.22 ± 11.83 (70-115) | 103.81 ± 8.03 (80-115) | .01 |
| Word list (46/42) | 8.72 ± 1.60 (4-10) | 9.26 ± 1.38 (4-10) | .09 |
| Working memory (46/42) | 6.46 ± 2.14 (4-13) | 7.48 ± 2.36 (3-14) | .04 |
| TMT A (45/41) | 36.67 ± 14.69 (16-80) | 26.12 ± 8.98 (10-50) | <.01* |
| TMT B (45/41) | 87.50 ± 50.49 (37-304) | 59.44 ± 21.51 (29-122) | <.01* |
| Processing speed (46/42) | 63.70 ± 19.10 (29-111) | 76.69 ± 11.32 (49-99) | <.01* |
| *Clinical measures* |  |  |  |
| PANSS Positive (46) | 20.83 ± 6.83 (7-39) | / | / |
| PANSS Negative (46) | 22.17 ± 7.64 (8-38) | / | / |
| PANSS GP (46) | 40.50 ± 11.41 (16-62) | / | / |
| PANSS Total (46) | 83.57 ± 22.31 (33-134) | / | / |
| SAPS Delusion (45) | 21.18 ± 10.70(0-48) | / | / |
| SANS Anhedonia (45) | 12.11 ± 4.97 (4-21) | / | / |
| CPZ (44) in mg | 380.27 ± 194.11 (33.25-1020) | / | / |

**S-Table 2.** Prior means and variances of parameters in computational models. α was estimated in logit-space, κ and *ω*_3_ were estimated in log space.

|  | Prior Mean | Prior Variance |
| --- | --- | --- |
| *Decision Model for all learning models* |  |  |
| *Softmax* |  |  |
| $\rho_{\text{rew}}$ | 1 | 1 |
| $\rho_{\text{pun}}$ | 0 | 1 |
| *Learning models* |  |  |
| *RL* |  |  |
| $\alpha$ | .5 | 1 |
| $\alpha_{\text{win}}$ | .6 | 1 |
| $\alpha_{\text{loss}}$ | .4 | 1 |
| *HGF* |  |  |
| $\omega$ (HGF2+3) | -1.5 | 0 |
| $\kappa$ (HGF3) | 1 | 1 |
| $\omega_3$ (HGF3) | -6 | 1 |
| $\mu_3^{(0)}$ (HGF3-V) | 0 | 1 |

**S-Table 3.** Explorative correlations (Spearman’s rank correlation coefficient) between model parameters that differed between groups and measures of cognitive functioning as well as symptom domains within patients fit better than chance by the model (n=31). * indicates significance surviving Bonferroni-correction for six cognitive tests. For completeness, we also report correlations between these two parameters and independent cognitive measures in HC.

| | $\mu_3^{(0)}$ | | $\kappa$ | |
| --- | --- | --- | --- | --- |
|  | r | p | r | p |
| *Cognitive measures (in PSZ)* |  |  |  |  |
| Verbal Intelligence (31) | -.35 | .06 | -.26 | .17 |
| Word list (31) | -.18 | .33 | -.15 | .43 |
| Working memory (31) | -.22 | .23 | -.25 | .18 |
| TMT A (30) | -.46 | .01 | -.33 | .08 |
| TMT B (30) | -.56* | <.001* | -.37 | .05 |
| Processing speed (31) | -.56* | <.001* | -.22 | .23 |
| *Clinical measures (in PSZ)* |  |  |  |  |
| PANSS Positive (31) | .25 | .17 | .26 | .16 |
| PANSS Negative (31) | .24 | .19 | .06 | .75 |
| PANSS GP (31) | .28 | .12 | .30 | .10 |
| PANSS Total (31) | .31 | .09 | .21 | .25 |
| SAPS Delusion (30) | .16 | .39 | .30 | .22 |
| SANS Anhedonia (30) | .32 | .09 | .07 | .72 |
| CPZ in mg (30) | .11 | .57 | .31 | .09 |
| *Cognitive measures (in HC)* |  |  |  |  |
| Verbal Intelligence (41) | .09 | .57 | .09 | .57 |
| Word list (41) | .08 | .60 | .17 | .29 |
| Working memory (41) | .04 | .80 | .11 | .48 |
| TMT A (40) | -.11 | .49 | -.09 | .57 |
| TMT B (40) | -.04 | .79 | .09 | .59 |
| Processing speed (41) | -.08 | .61 | -.16 | .33 |

**S-Table 4.**
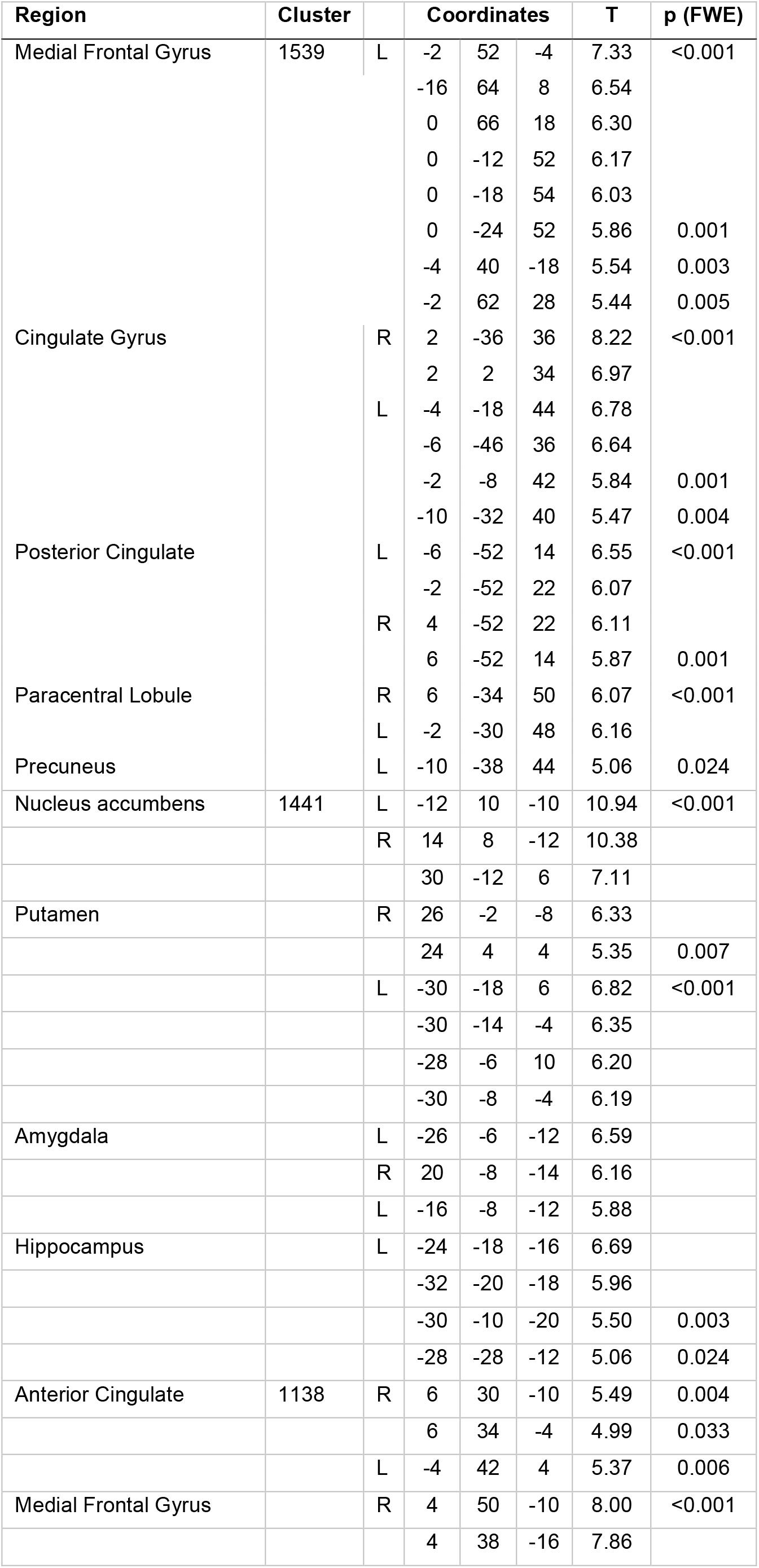
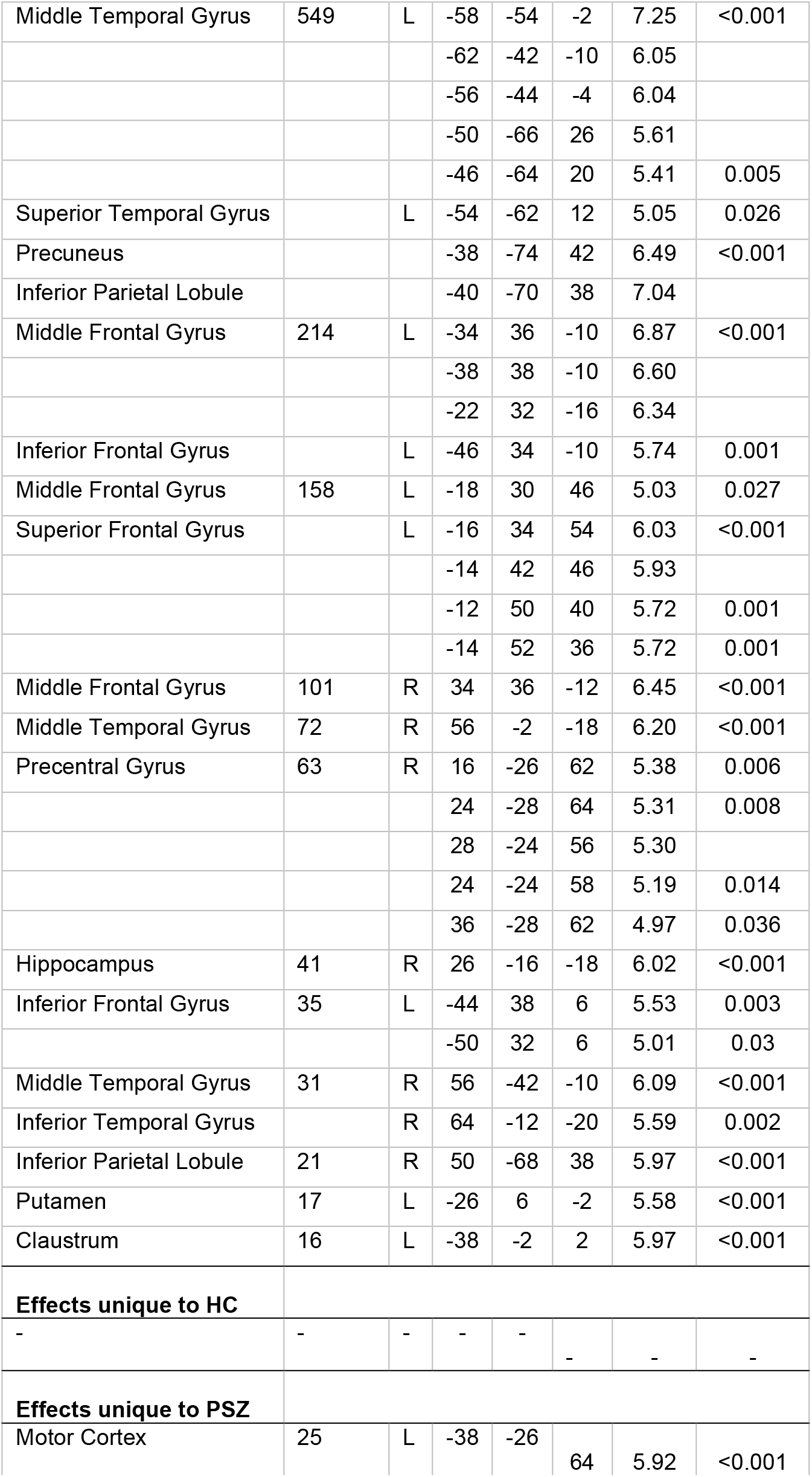
ε2 related activation in all participants, p-FWE_wholebrain_<0.05, k=10

**S-Table 5.** ε3 related activation in all participants, p-FWE_wholebrain_<0.05, k=10

| Region | Cluster |  | Coordinates |  |  | T | p (FWE) |
| --- | --- | --- | --- | --- | --- | --- | --- |
| Inferior Frontal Gyrus | 84 | R | 44 | 12 | 26 | 6.44 | <0.001 |
| Inferior Parietal Lobule | 28 | L | -42 | -38 | 40 | 5.69 | 0.001 |
| Middle Frontal Gyrus | 26 | L | -30 | 2 | 60 | 5.48 | 0.004 |
| Precuneus | 20 | L | -32 | -50 | 50 | 5.17 | 0.015 |
| Inferior Frontal Gyrus | 19 | L | -50 | 4 | 36 | 5.30 | 0.009 |
| Precuneus | 7 | R | 10 | -72 | 44 | 5.30 | 0.009 |
| Insula | 5 | L | -40 | 8 | 24 | 5.46 | 0.004 |
| Superior Frontal Gyrus | 4 | L | -4 | 10 | 54 | 5.03 | 0.027 |
| Precuneus | 3 | L | -30 | -74 | 28 | 5.22 | 0.012 |
| Putamen | 2 | L | -14 | 10 | -12 | 4.96 | 0.036 |
| Inferior Parietal Lobule | 2 | L | -40 | -48 | 44 | 4.92 | 0.044 |
| **Effects unique to HC** |  |  |  |  |  |  |  |
| - | - | - | - | - | - | - | - |
| **Effects unique to PSZ** |  |  |  |  |  |  |  |
| - | - | - | - | - | - | - | - |

**S-Table 6.** Ψ2 related activation in all participants, p-FWE_wholebrain_<0.05, k=10

| Region | Cluster |  | Coordinates |  |  | T | p (FWE) |
| --- | --- | --- | --- | --- | --- | --- | --- |
| Cingulate Gyrus | 564 |  | 6 | 28 | 38 | 7.77 | <0.001 |
|  |  |  | 10 | 18 | 42 | 7.32 |  |
|  |  | L | -6 | 24 | 38 | 6.27 |  |
| Anterior Cingulate |  | R | 4 | 38 | 28 | 5.34 | 0.007 |
| Inferior Frontal Gyrus | 348 | L | -28 | 24 | -6 | 7.87 | <0.001 |
| Clastrum |  |  | -30 | 20 | 8 | 6.61 |  |
|  |  |  | -32 | 18 | 2 | 6.46 |  |
| Insula |  |  | -40 | 16 | 8 | 6.09 |  |
| Inferior Frontal Gyrus | 332 | R | 30 | 24 | -8 | 8.45 | <0.001 |
| Middle Frontal Gyrus | 180 | R | 36 | 40 | 24 | 6.01 | <0.001 |
|  |  |  | 42 | 30 | 30 | 5.91 | 0.001 |
|  |  |  | 36 | 24 | 34 | 5.57 | 0.002 |
|  |  |  | 36 | 50 | 18 | 5.12 | 0.019 |
| Inferior Parietal Lobule | 125 | R | 52 | -46 | 38 | 6.26 | <0.001 |
|  |  |  | 44 | -36 | 40 | 5.51 | 0.003 |
| Inferior Parietal Lobule | 118 | L | -42 | -58 | 52 | 5.51 | 0.003 |
|  |  |  | -42 | -52 | 44 | 5.45 | 0.004 |
|  |  |  | -46 | -52 | 54 | 5.33 | 0.008 |
|  |  |  | -48 | -50 | 50 | 5.07 | 0.024 |
| Middle Frontal Gyrus | 111 | L | -38 | 56 | 12 | 6.65 | <0.001 |
|  |  |  | -40 | 54 | -2 | 5.85 | 0.001 |
|  |  |  | -46 | 48 | -8 | 5.15 | 0.016 |
| Cerebellum | 109 | L | -32 | -58 | -30 | 7.36 | <0.001 |
| Cerebellum | 92 | R | 36 | -62 | -28 | 7.24 | <0.001 |
|  |  |  | 46 | -66 | -26 | 5.44 | 0.005 |
| Middle Frontal Gyrus | 86 | L | -36 | 28 | 30 | 5.77 | 0.001 |
| Inferior Frontal Gyrus | 53 | L | -36 | 4 | 32 | 6.01 | <0.001 |
| Middle Frontal Gyrus |  |  | -42 | 10 | 36 | 5.49 | 0.004 |
| Red Nucleus | 52 | L | -6 | -24 | -4 | 7.47 | <0.001 |
| Globus Pallidus | 41 | L | -14 | 2 | -2 | 6.03 | <0.001 |
| Thalamus |  |  | -12 | -10 | 0 | 5.29 | 0.009 |
| Inferior Parietal Lobule | 40 | L | -46 | -38 | 42 | 6.32 | <0.001 |
| Thalamus | 29 | R | 12 | -6 | 8 | 5.68 | 0.001 |
| Precentral Gyrus | 24 | R | 44 | 2 | 38 | 5.21 | 0.013 |
| Superior Frontal Gyrus | 27 | L | -6 | 12 | 54 | 5.78 | 0.001 |
| Middle Frontal Gyrus |  |  | -34 | -2 | 48 | 5.67 | 0.002 |
| Cingulate Gyrus |  |  | -4 | 14 | 46 | 5.01 | 0.030 |
| Caudate | 20 | R | 14 | 12 | 8 | 5.80 | 0.001 |
| Inferior Parietal Gyrus | 14 | L | -32 | -48 | 38 | 5.2 | 0.013 |
| Inferior Frontal Gyrus | 11 | R | 48 | 22 | 10 | 5.53 | 0.003 |
| Middle Temporal Gyrus | 11 | L | -56 | -30 | -10 | 5.25 | 0.010 |
| Insula | 10 | R | 44 | 16 | 4 | 5.54 | 0.003 |
| **Effects unique to HC** |  |  |  |  |  |  |  |
| - | - | - | - | - | - | - | - |
| **Effects unique to PSZ** |  |  |  |  |  |  |  |
| - | - | - | - | - | - | - | - |

**S-Table 7.** Ψ3 related activation in all participants, p-FWE_wholebrain_<0.05, k=10

| Region | Cluster |  | Coordinates |  |  | T | p (FWE) |
| --- | --- | --- | --- | --- | --- | --- | --- |
| Medial Frontal Gyrus | 126 | R | 6 | 26 | 40 | 6.01 | <0.001 |
| Cingulate Gyrus |  | R | 10 | 18 | 42 | 6.15 |  |
|  |  |  | 8 | 12 | 46 | 5.89 | 0.001 |
| Insula |  | L | -32 | 20 | 10 | 5.57 | 0.002 |
|  |  |  | -36 | 18 | 10 | 5.52 | 0.003 |
|  |  | R | 34 | 22 | 6 | 5.56 |  |
| Clastrum |  | L | -28 | 24 | 4 | 5.49 | 0.004 |
| Inferior Frontal Gyrus | 104 | R | 30 | 24 | -8 | 6.34 | <0.001 |
| Inferior Frontal Gyrus | 97 | L | -28 | 24 | -6 | 5.90 | 0.001 |
| Clastrum | 86 | R | 36 | -62 | -28 | 6.53 | <0.001 |
|  |  |  | 32 | -68 | -22 | 5.65 | 0.002 |
|  | 44 | L | -32 | -56 | -30 | 6.28 | <0.001 |
| Red nucleus | 12 | L | -6 | -24 | -4 | 5.91 | <0.001 |
| Inferior Frontal Gyrus | 11 | L | -36 | 4 | 32 | 5.35 | 0.007 |
| Anterior Cingulate | 11 | R | 8 | 28 | 30 | 5.39 | 0.006 |
|  |  | L | -6 | 30 | 30 | 5.36 |  |
| Cingulate Gyrus |  |  | -8 | 22 | 38 | 5.34 | 0.007 |
| **Effects unique to HC** |  |  |  |  |  |  |  |
| - | - | - | - | - | - | - | - |
| **Effects unique to PSZ** |  |  |  |  |  |  |  |
| Cerebellum | 13 | R | 36 | -64 | -22 | 5.25 | 0.007 |
|  |  |  | 38 | -56 | -26 | 5.2 | 0.009 |
|  | 11 | L | -38 | -58 | -30 | 5.1 | 0.012 |

**S-Table 8.**
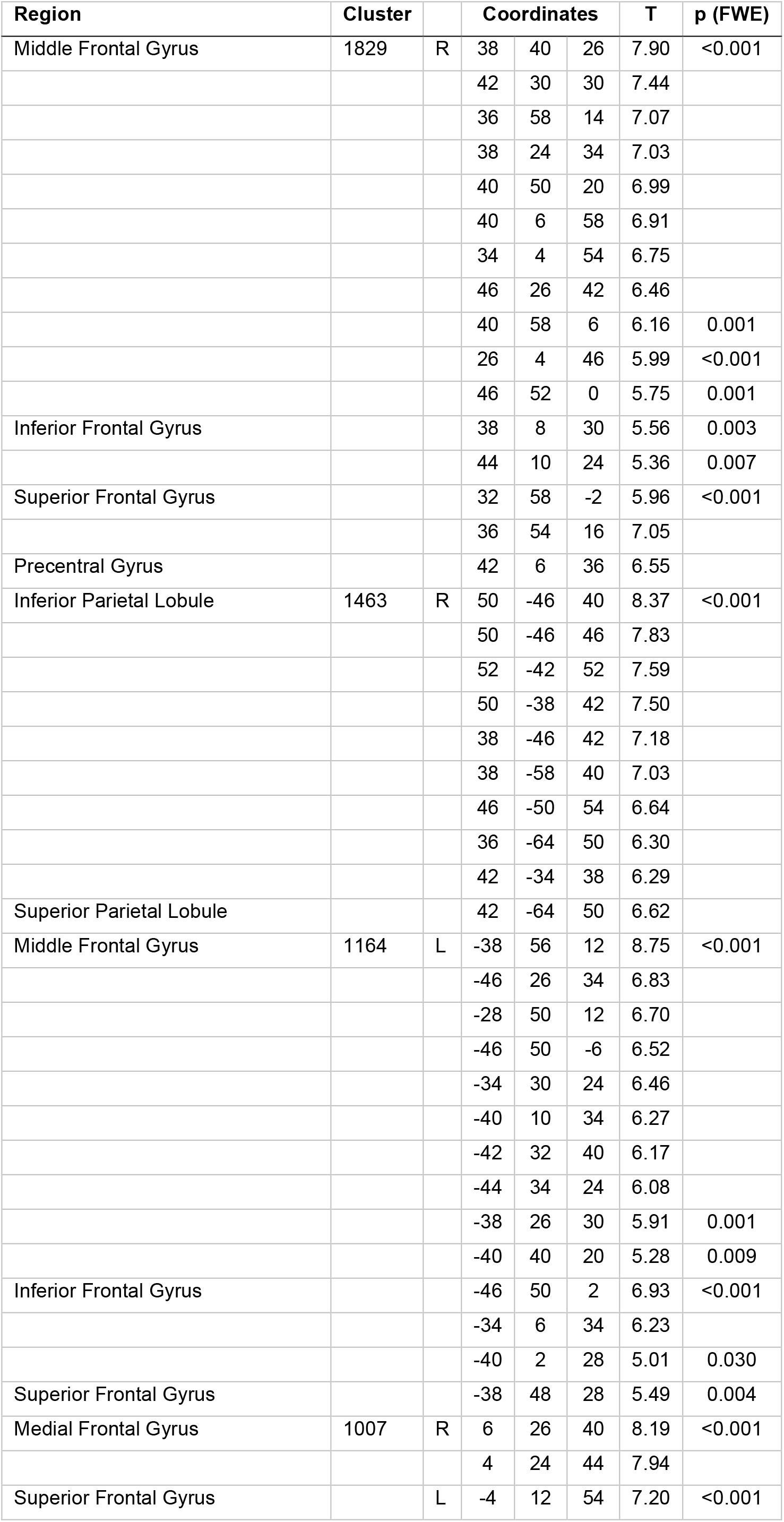
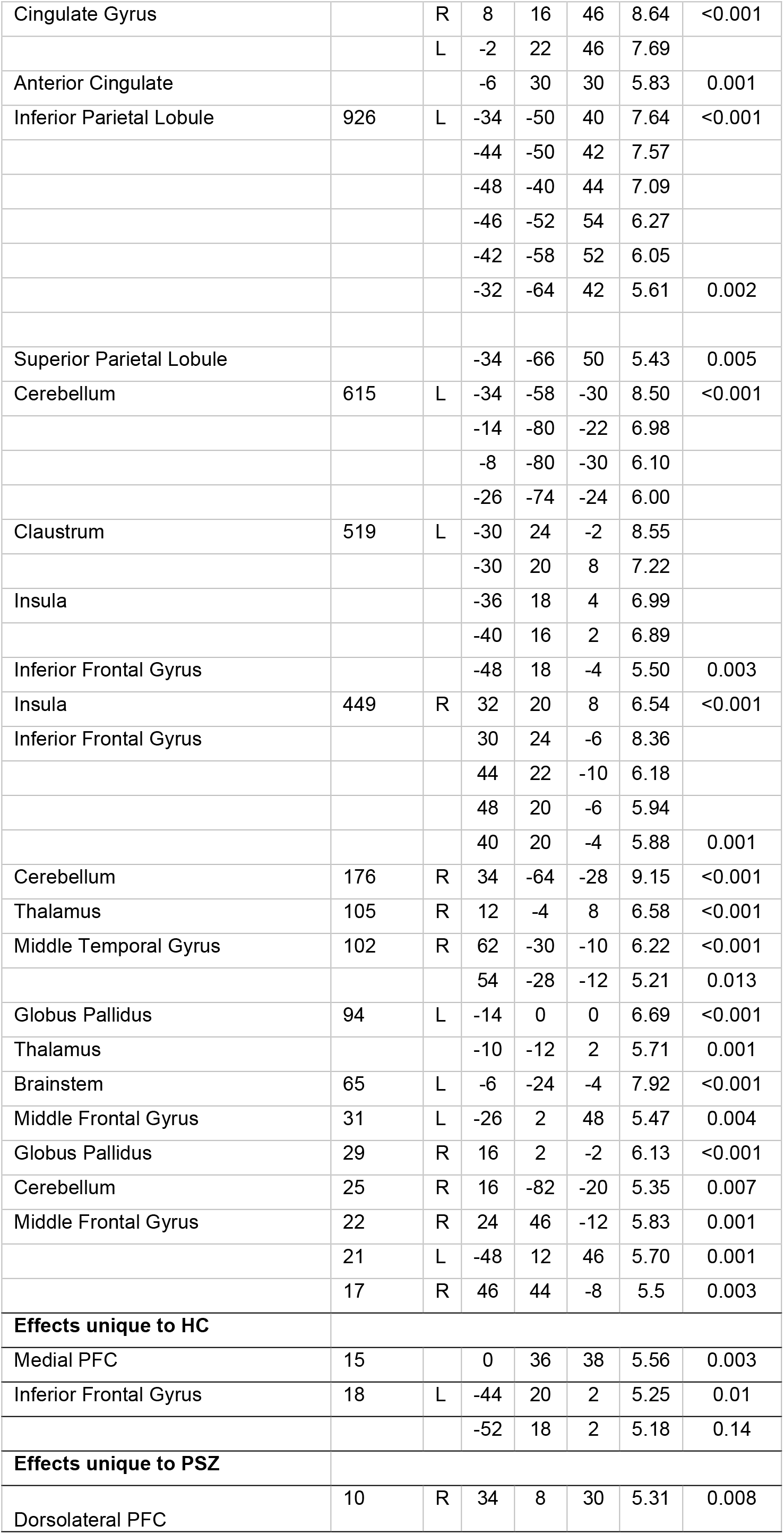
µ3 related activation in all participants, p-FWE_wholebrain_<0.05, k=10

**S-Table 9.** Reward prediction error related activation (RPE δ from reinforcement learning model), over all participants, FWE corrected p < 0.05, k = cluster size (>=10).

| Region | Cluster |  | Coordinates |  |  | T | p (FWE) |
| --- | --- | --- | --- | --- | --- | --- | --- |
| Superior Frontal Gyrus | 3908 | L | -16 | 36 | 52 | 9.07 | <0.001 |
|  |  |  | -14 | 54 | 30 | 7.84 |  |
|  |  | R | 14 | 54 | 36 | 7.32 |  |
|  |  |  | 18 | 36 | 54 | 7.31 |  |
| Medial Frontal Gyrus |  | L | -2 | 64 | 8 | 8.61 |  |
|  |  |  | -4 | 58 | 4 | 8.34 |  |
|  |  |  | -8 | 56 | -12 | 7.36 |  |
|  |  | R | 2 | 66 | 20 | 8.54 |  |
|  |  |  | 6 | 56 | -4 | 8.40 |  |
|  |  |  | 2 | 64 | 24 | 8.01 |  |
|  |  |  | 2 | 42 | -14 | 7.96 |  |
|  |  |  | 4 | 36 | -12 | 7.74 |  |
|  |  |  | 2 | 62 | 28 | 7.64 |  |
| Middle Frontal Gyrus |  |  | 30 | 34 | -14 | 7.33 |  |
| Anterior Cingulate |  | L | -6 | 40 | -8 | 8.40 |  |
|  |  | R | 4 | 40 | -8 | 7.72 |  |
| Cingulate |  | L | -2 | -48 | 32 | 7.81 |  |
|  |  |  | -6 | -46 | 34 | 7.74 |  |
|  |  |  | -10 | -48 | 36 | 7.25 |  |
|  |  |  | -10 | -32 | 40 | 6.10 | 0.002 |
|  |  |  | 0 | -2 | 36 | 6.05 |  |
|  |  |  | -4 | -8 | 44 | 5.92 | 0.004 |
|  |  | R | 2 | -34 | 36 | 9.29 | <0.001 |
|  |  |  | 2 | -10 | 38 | 6.67 |  |
|  |  |  | 4 | -14 | 36 | 6.35 | 0.001 |
| Posterior Cingulate |  |  | -8 | -56 | 16 | 8.66 | <0.001 |
|  | 1845 | L | -2 | -52 | 22 | 9.50 | <0.001 |
| Paracentral Lobule |  | R | 6 | -34 | 48 | 5.39 | 0.025 |
| Putamen | 1218 | L | -14 | 8 | -10 | 9.56 | <0.001 |
|  |  |  | -28 | -6 | 4 | 9.14 |  |
|  |  |  | -30 | -6 | -6 | 8.19 |  |
|  |  |  | -26 | 8 | 2 | 6.71 |  |
| Nucleus accumbens |  | L | -8 | 12 | -10 | 9.25 |  |
| Amygdala |  |  | -26 | -8 | -18 | 9.47 |  |
| Hippocampus |  |  | -28 | -20 | -18 | 8.42 |  |
| Clastrum |  |  | -34 | -6 | -2 | 8.64 |  |
|  |  |  | -38 | 0 | 4 | 6.85 |  |
| Globus Pallidus |  |  | -14 | -8 | -10 | 6.16 | 0.001 |
| Middle Frontal Gyrus | 1039 | L | -42 | 38 | 2 | 8.42 | <0.001 |
|  |  |  | -36 | 38 | -10 | 7.46 |  |
|  |  |  | -40 | 34 | -14 | 7.33 |  |
|  |  |  | -42 | 50 | -4 | 6.98 |  |
|  |  |  | -46 | 44 | -4 | 6.84 |  |
|  |  |  | -22 | 34 | -14 | 6.64 |  |
| Inferior Frontal Gyrus |  |  | -50 | 38 | 8 | 7.54 |  |
|  |  |  | -46 | 38 | -10 | 7.20 |  |
|  |  |  | -40 | 26 | -16 | 6.94 |  |
|  |  |  | -26 | 30 | -16 | 6.71 |  |
|  |  |  | -50 | 32 | -8 | 6.41 | 0.001 |
|  |  |  | -54 | 32 | 0 | 6.40 |  |
|  |  |  | -52 | 30 | 8 | 6.37 |  |
|  |  |  | -46 | 48 | 8 | 5.93 | 0.003 |
| Putamen | 1013 | R | 30 | -14 | 2 | 8.78 | <0.001 |
|  |  |  | 26 | 2 | 4 | 8.09 |  |
|  |  |  | 28 | -4 | 0 | 7.82 |  |
|  |  |  | 30 | -2 | -10 | 6.03 | 0.002 |
| Nucleus accumbens |  |  | 12 | 10 | -10 | 9.89 | <0.001 |
| Amygdala |  |  | 24 | -6 | -18 | 6.67 |  |
|  |  |  | 26 | -6 | -26 | 6.21 | 0.001 |
| Hippocampus |  |  | 30 | -12 | -20 | 7.40 | <0.001 |
|  |  |  | 28 | -20 | -16 | 7.00 |  |
| Clastrum |  |  | 34 | 2 | 2 | 6.43 | 0.001 |
| Middle Temporal Gyrus | 769 | L | -50 | -66 | 26 | 8.31 | <0.001 |
| Angular Gyrus |  |  | -52 | -68 | 30 | 8.26 |  |
| Precuneus |  |  | -40 | -68 | 36 | 7.77 |  |
| Superior Parietal Lobule |  |  | -42 | -64 | 50 | 6.20 | 0.001 |
|  |  |  | -36 | -74 | 48 | 6.09 | 0.002 |
| Cerebellum | 502 | L | -40 | -76 | -32 | 9.35 | <0.001 |
|  |  |  | -36 | -80 | -28 | 7.99 |  |
|  |  |  | -46 | -70 | -32 | 7.68 |  |
|  |  |  | -14 | -86 | -28 | 7.60 |  |
|  |  |  | -26 | -76 | -30 | 7.08 |  |
|  |  |  | -24 | -80 | -28 | 7.01 |  |
|  |  |  | -28 | -82 | -24 | 6.62 |  |
|  |  |  | -6 | -84 | -28 | 6.62 |  |
|  |  |  | -32 | -70 | -34 | 6.41 | 0.001 |
|  |  |  | -18 | -72 | -32 | 6.03 | 0.002 |
|  |  |  | -34 | -64 | -34 | 5.31 | 0.033 |
| Inferior Temporal Gyrus | 387 | L | -56 | -16 | -18 | 7.21 | <0.001 |
|  |  |  | -62 | -10 | -20 | 6.97 |  |
|  |  |  | -62 | -16 | -20 | 6.84 |  |
|  |  |  | -60 | -54 | -8 | 6.02 | 0.002 |
| Middle Temporal Gyrus | 387 | L | -64 | -48 | -4 | 7.05 | <0.001 |
|  |  |  | -60 | -34 | -10 | 7.02 |  |
|  |  |  | -54 | -36 | -14 | 5.91 | 0.004 |
|  |  |  | -58 | -50 | -12 | 5.74 | 0.007 |
| Inferior Parietal Lobule | 318 | R | 54 | -60 | 42 | 6.61 | <0.001 |
|  |  |  | 52 | -66 | 36 | 6.50 |  |
|  |  |  | 46 | -66 | 40 | 6.15 | 0.001 |
|  |  |  | 44 | -72 | 42 | 6.09 | 0.002 |
|  |  |  | 44 | -68 | 46 | 6.03 |  |
| Superior Parietal Lobule |  |  | 42 | -64 | 50 | 5.82 | 0.005 |
|  |  |  | 50 | -58 | 26 | 5.36 | 0.028 |
|  |  |  | 46 | -60 | 28 | 5.30 | 0.034 |
|  |  |  | 54 | -60 | 24 | 5.29 | 0.036 |
| Cerebellum | 160 | R | 20 | -42 | -24 | 7.02 | <0.001 |
|  |  |  | 16 | -46 | -22 | 6.91 |  |
|  |  |  | 22 | -54 | -24 | 6.57 |  |
|  |  |  | 28 | -48 | -26 | 6.05 | 0.002 |
| Middle Temporal Gyrus | 144 | R | 62 | -38 | -14 | 7.07 | <0.001 |
|  |  |  | 64 | -30 | -14 | 6.75 |  |
|  |  |  | 64 | -42 | -4 | 6.32 | 0.001 |
|  |  |  | 58 | -42 | -12 | 6.13 | 0.002 |
| Cerebellum | 113 | R | 26 | -82 | -22 | 6.53 | <0.001 |
|  |  |  | 20 | -74 | -22 | 5.81 | 0.005 |
|  |  |  | 22 | -86 | -28 | 5.70 | 0.008 |
|  |  |  | 10 | -84 | -26 | 5.38 | 0.026 |
| Thalamus | 89 | R | 14 | -6 | 14 | 6.44 | 0.001 |
| Caudate |  |  | 18 | -2 | 22 | 6.25 |  |
|  | 69 | L | -14 | -10 | 18 | 6.36 | 0.001 |
| Thalamus |  |  | -8 | -4 | 16 | 5.97 | 0.003 |
| Middle Frontal Gyrus | 57 |  | 34 | 30 | 52 | 6.51 | <0.001 |
| Inferior Temporal Gyrus | 44 | R | 50 | -8 | -22 | 5.73 | 0.007 |
|  |  |  | 60 | -14 | -22 | 5.71 | 0.008 |
| Middle Temporal Gyrus |  |  | 62 | -8 | -14 | 6.01 | 0.003 |
|  |  |  | 56 | -20 | -20 | 5.39 | 0.025 |
| Medial Frontal Gyrus | 39 | L | 0 | -12 | 52 | 6.10 | 0.002 |
| Cerebellum | 30 | R | 44 | -74 | -28 | 6.98 | <0.001 |
|  | 22 | L | -22 | -56 | -20 | 6.14 | 0.002 |
|  | 17 | L | -26 | -42 | -30 | 6.2 | 0.001 |
| Superior Parietal Lobule | 14 |  | -26 | -40 | 56 | 6.03 | 0.002 |
| Culmen | 13 | L | -18 | -38 | -24 | 5.57 | 0.013 |
| Insula | 11 | R | 36 | 2 | 22 | 6.29 | 0.001 |
| Inferior Parietal Lobule | 10 | L | -64 | -36 | 24 | 5.65 | 0.010 |

**S-Table 10.** Small volume corrections for region of interests (ROI) in midbrain and ventral striatum. Bilateral ROIs were used for midbrain as in Iglesias et al. 2016 based on a study by Bunzeck and Düzel (2006) and for ventral striatum as in Schlagenhauf et al. (2014)^8, 16, 17^.

| Region |  |  | Coordinates |  |  | T/F | p (FWE) |
| --- | --- | --- | --- | --- | --- | --- | --- |
| Second level precision-weighted prediction error $\epsilon_2$ | | | | | | | |
| Midbrain ROI | All HC and SZ, T | L | -8 | -18 | -10 | 3.94 | .01 |
|  | HC vs SZ, F | R | 12 | -18 | -14 | 12.97 | .08 |
| Ventral striatum ROI | All HC and SZ, T | FWE<sub>wholebrain</sub> < .05, see S-Table 3 |  |  |  |  |  |
|  | HC vs SZ, F | R | 16 | 2 | -6 | 6.23 | .35 |
| Reward prediction error from reinforcement learning model $\delta$ | | | | | | | |
| Midbrian ROI | All HC and SZ, T | L | -6 | -14 | -10 | 4.03 | .02 |
|  | HC vs SZ, F | R | 14 | -18 | -12 | 11.60 | .17 |
| Ventral striatum ROI | All HC and SZ, T | FWE<sub>wholebrain</sub> < .05, see S-Table 8 |  |  |  |  |  |
|  | HC vs SZ, F | L | -14 | 4 | -4 | 9.49 | .12 |

**S-Figure 1.**
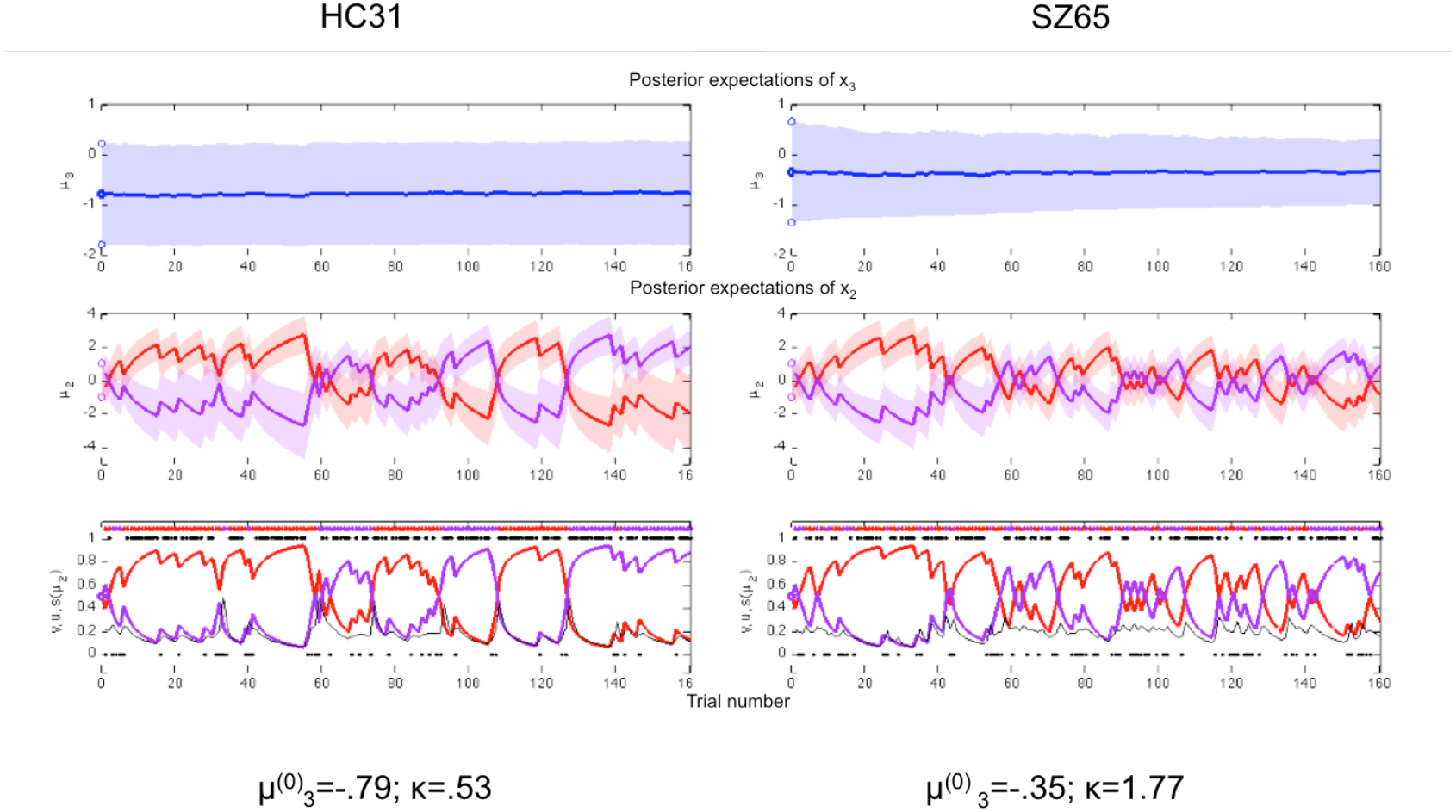
Example trajectories of one healthy control and one patient with the initial belief over mu3 and kappa close to the group mean illustrating the differences in learning on the second and the third level of the HGF.

**S-Figure 2.**
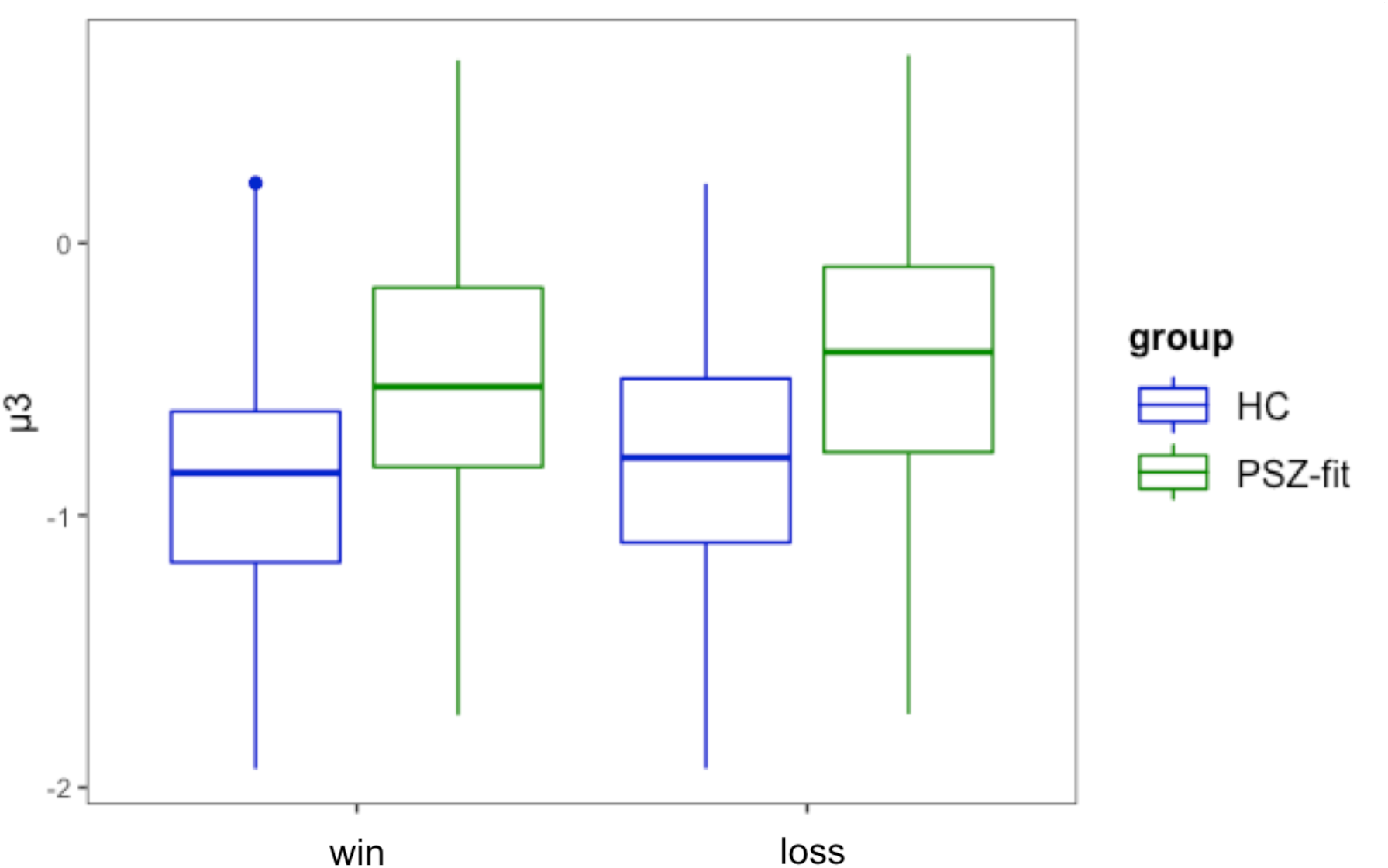
Analysis of trial-wise posterior means 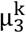 of inferred environmental volatility in a mixed-effects regression model with group and feedback as predictors revealed a significant main effect of group (t=3.22, p=0.01) due to higher estimates of volatility in PSZ and a main effect of feedback (t=16.124, p<.001) due to higher estimates of volatility after losses compared to rewards. A group x feedback interaction (t=2.50, 0.02) was also significant and driven by a larger difference in environmental volatility after having received losses versus rewards in PSZ as compared to HC (S-Figure 2).

**S-Figure 3.**
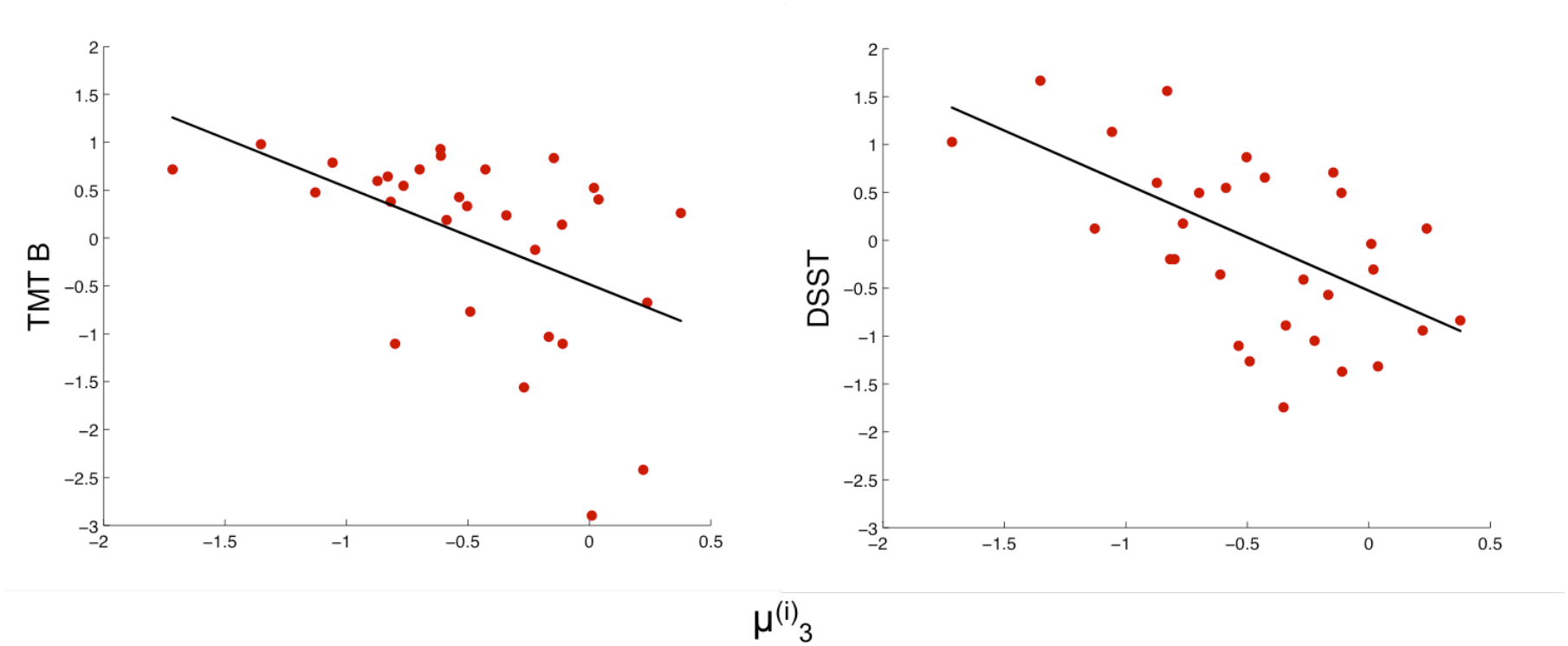
Bonferroni-corrected significant correlations between 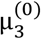 and z-scored measures of executive functioning (TMT B) and cognitive speed (DSST) in patients fit better than chance by the model (n=31), Spearman rank correlation coefficient -.56, p<.001 for TMT B and -.56, p<.001 for DSST.

**S-Figure 4.**
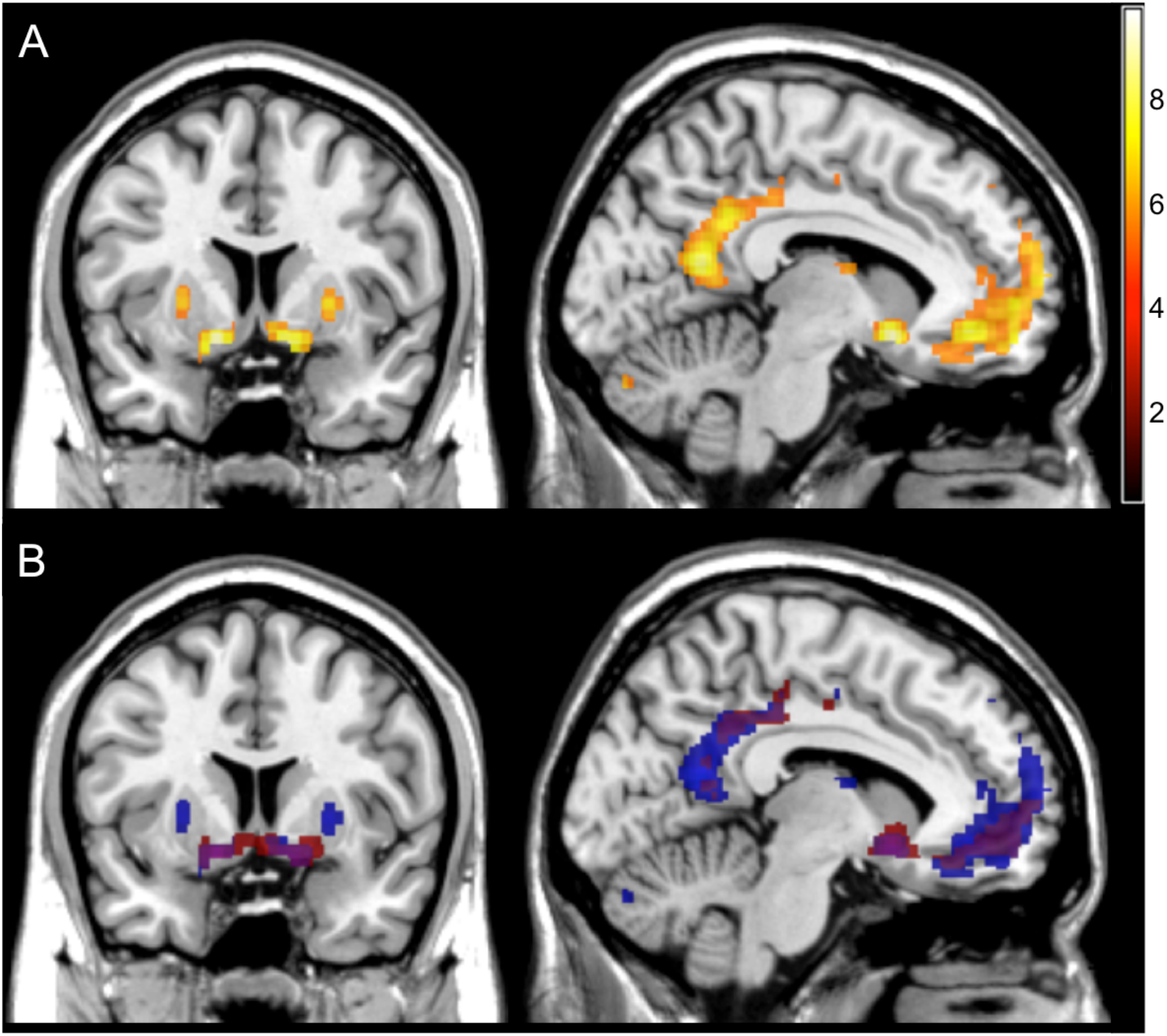
A) PE-related signals based on RL model, p-FWE_wholebrain_<0.05, k=10, [7 −8]; B) Overlay of RRE from RL model in blue and second-level precision weighted RPE from HGF model in red, p-FWE_wholebrain_<0.05, k=10, [7 −8];

**S-Figure 5.**
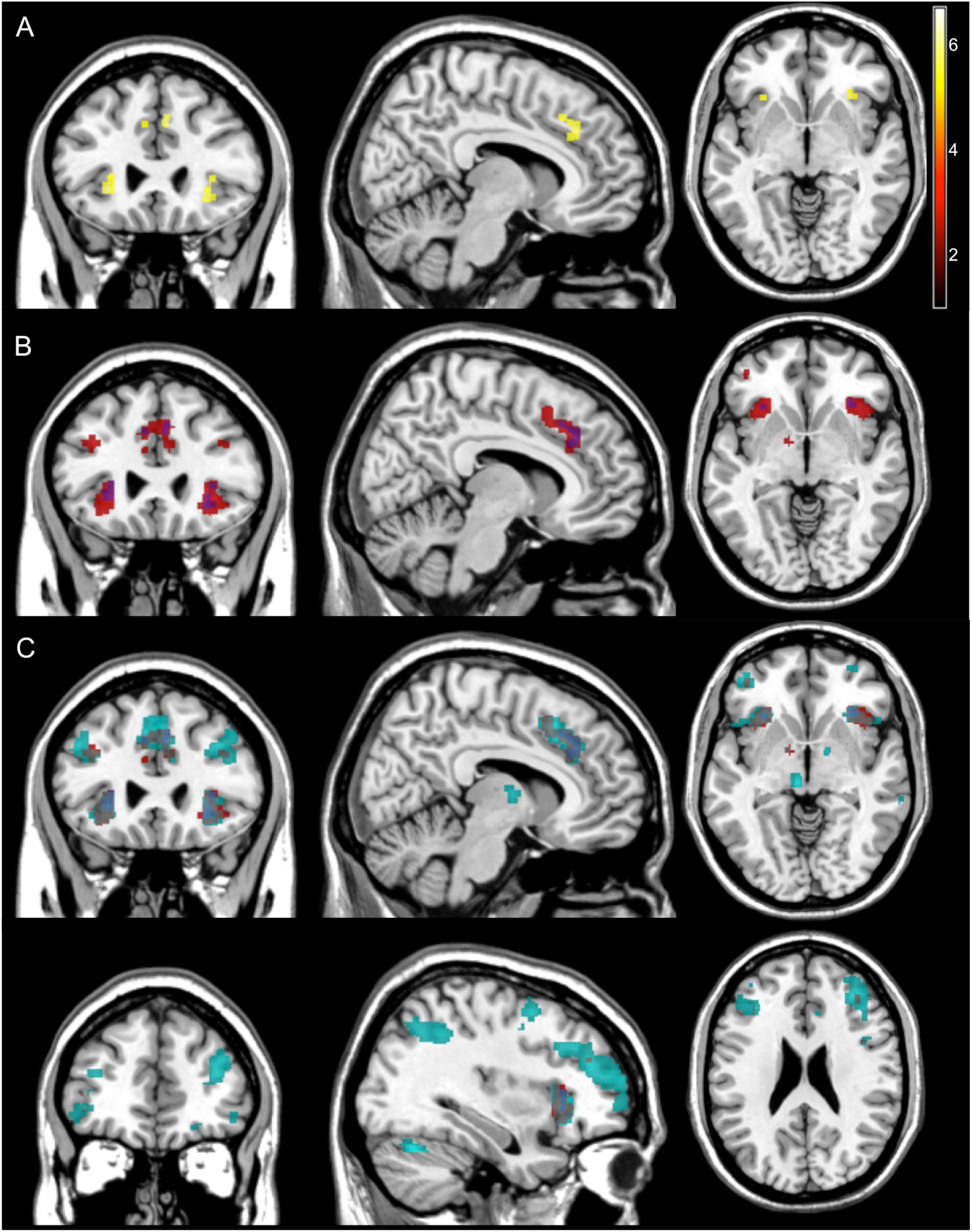
A) BOLD signal related to third-level precision, p-FWE_wholebrain_<0.05, k=10, [8 26 −4]; B) overlay of second-level (red) and third-level precision (blue), p-FWE_wholebrain_<0.05, k=10; C) overlay of second-level (red) and third-level precision (blue), environmental volatility (turqoise) at [8 26 −4] and [34 44 24], p-FWE_wholebrain_<0.05, k=10;

